# Endogenous Retroviral Elements Generate Pathologic Neutrophils and Elastase Rich Exosomes in Pulmonary Arterial Hypertension

**DOI:** 10.1101/2021.01.08.426001

**Authors:** Shalina Taylor, Kévin Contrepois, Bérénice A. Benayoun, Lihua Jiang, Sarasa Isobe, Lingli Wang, Stavros Melemenidis, Mehmet O. Ozen, Shoichiro Otsuki, Tsutomu Shinohara, Andrew J. Sweatt, Jordan Kaplan, Aiqin Cao, Jan-Renier Moonen, David P. Marciano, Mingxia Gu, Kazuya Miyagawa, Patricia A. Del Rosario, Andrew Hsi, A. A. Roger Thompson, Maria-Eugenia Ariza, Utkan Demirci, Roham T. Zamanian, Francois Haddad, Mark R. Nicolls, Michael P. Snyder, Marlene Rabinovitch

## Abstract

Neutrophil elastase (NE) is implicated in pulmonary arterial hypertension (PAH) but the role of neutrophils in the pathogenesis of PAH is unclear. Here we show that neutrophils from PAH vs. control subjects produce and release increased NE associated with enhanced extracellular trap formation. PAH neutrophils are highly adherent and show decreased migration consistent with increased vinculin, identified on proteomic analysis and previously linked to an antiviral response. This was substantiated by a transcriptomic interferon signature in PAH neutrophils and an increase in human endogenous retrovirus (HERV-K) envelope protein. NE and interferon genes are induced by HERV-K envelope and vinculin is increased by HERV-K dUTPase that is elevated in PAH plasma. Neutrophil exosomes from PAH plasma contain increased NE and HERV-K envelope and induce pulmonary hypertension in mice, that is mitigated by the NE inhibitor and antiviral agent, elafin. Thus, elevated HERVs explain pathological neutrophils linked to PAH induction and progression.

## INTRODUCTION

Pulmonary arterial hypertension (PAH) is a disease characterized by occlusion of distal pulmonary arteries resulting in increased resistance to flow and culminating in heart failure. Current treatments improve quality of life and survival primarily by dilating pulmonary arteries, but these agents do not address the mechanism underlying the progressive pulmonary vascular pathology (reviewed in ^1^). The appearance of fragmented elastic laminae in pulmonary arteries from patients with PAH suggested the presence of heightened elastase activity in the vessel wall^2^. Increased pulmonary arterial smooth muscle cell (SMC) elastase activity was identified as neutrophil elastase (NE) in human cells and in experimental animals with pulmonary hypertension^3^ Multiple sequelae of heightened NE activity are linked to the pathogenesis of PAH. Degradation of elastin increases vascular stiffness^4^ and elastin peptides are highly chemotactic to monocytes^5^ leading to chronic perivascular inflammation. NE induces release of growth factors from the extracellular matrix^6^ and can activate growth factor receptors^7^ resulting in SMC proliferation^8^. Moreover, inhibition of NE with synthetic inhibitors^9^ or with the recombinant human elastase inhibitor elafin^10^, is effective in preventing and in reversing pulmonary hypertension in animal models and pulmonary arterial pathology in human organ cultures^10^.

Neutrophils are major sources of NE; increased release of NE is observed in PAH neutrophils^11^, and NE levels are elevated in plasma from PAH patients^12^. Morevoer, the neutrophil to lymphocyte ratio is increased in PAH and correlates with clinical deterioration judged by New York Heart Association (NYHA) functional class, and event free survival^13^. Neutrophils are recruited to sites of tissue injury and play a major role in regulating the innate immune response against invading pathogens and the adaptive immune response through interactions with T and B cells. In addition to the release of NE and other antimicrobial proteins, neutrophil functions such as extracellular trap formation (NETs)^14^, adhesion and migration are important in response to injury^15^. These functions are inter-related, i.e., reducing NE is associated with decreased neutrophil adhesion and migration, as was shown with NE inhibitors^16^ or in NE deficient mice^17^.

NE is critical in the process of neutrophil chromatin release causing NETs, and NE associated with NETs remains proteolytically active and can cause tissue damage^18^. Elevated markers of NET formation have been identified in end-stage plexiform lesions in patients with PAH^12^. In addition to local effects, NE can have remote consequences by being packaged in exosomes. Bronchial lavage from COPD patients contains exosomes enriched with NE that caused alveolar destruction when injected intratracheally in mice^19^.

To understand their contribution to the pathogenesis of PAH, we isolated circulating neutrophils from PAH patients vs. healthy controls and assessed the functions of these cells. We then carried out proteomic and transcriptomic profiling to establish the mechanism underlying the abnormalities observed. Our results indicated an antiviral response in PAH neutrophils that we attributed to an elevation in human endogenous retroviral HERV-K envelope protein. Neutrophil HERV-K envelope and increased circulating HERV-K dUTPase protein explained increased adhesion, reduced migration and heightened NE production/NET formation observed in PAH neutrophils. PAH neutrophil exosomes contain increased NE and HERV-K, and induced pulmonary hypertension in mice, except when pretreated with elafin, a NE inhibitor with antiviral properties^20^. Taken together, our studies can explain how neutrophil dysfunction contributes to PAH and can be targeted therapeutically.

## RESULTS

### Increased NE protein and NE-mediated NETosis in PAH neutrophils

Neutrophils isolated from the blood of PAH patients and control subjects were confirmed to be >95% pure as described in the *Methods* section and shown in **Supplementary Figure 1a**. Demographics and other data related to the cohorts used in each of the experiments are found in **Supplementary Tables 1-4**. While we requested samples from patients with idiopathic PAH, a small proportion (3/68) were later reclassified as having drug and toxin related Group 1 PAH^21^, and 5/68 had some cardiopulmonary co-morbidities not thought to be causing the PAH. We observed a greater than two-fold increase in levels of NE protein in PAH neutrophils when compared to those of control subjects as assessed by western immunoblot (**Figure 1a**). A comparable increase in NE activity was established using a fluorescent labeled (DQ) elastin substrate (**Figure 1b**). We also confirmed increased release of NE activity following interleukin-8 (IL-8) stimulation of PAH vs. control neutrophils (**Figure 1c**). To confirm selective digestion of DQ elastin by NE, we used an inhibitor of NE, N-methoxysuccinyl-Ala-Ala-Pro-Val-chloromethyl ketone. While more than 2/3 of elastase activity measured by DQ elastin could be attributed to NE, there also appeared to be an elevation in non-NE elastolytic activity in PAH neutrophils (**Supplementary Figure 1b and 1c**).

**Figure 1.**
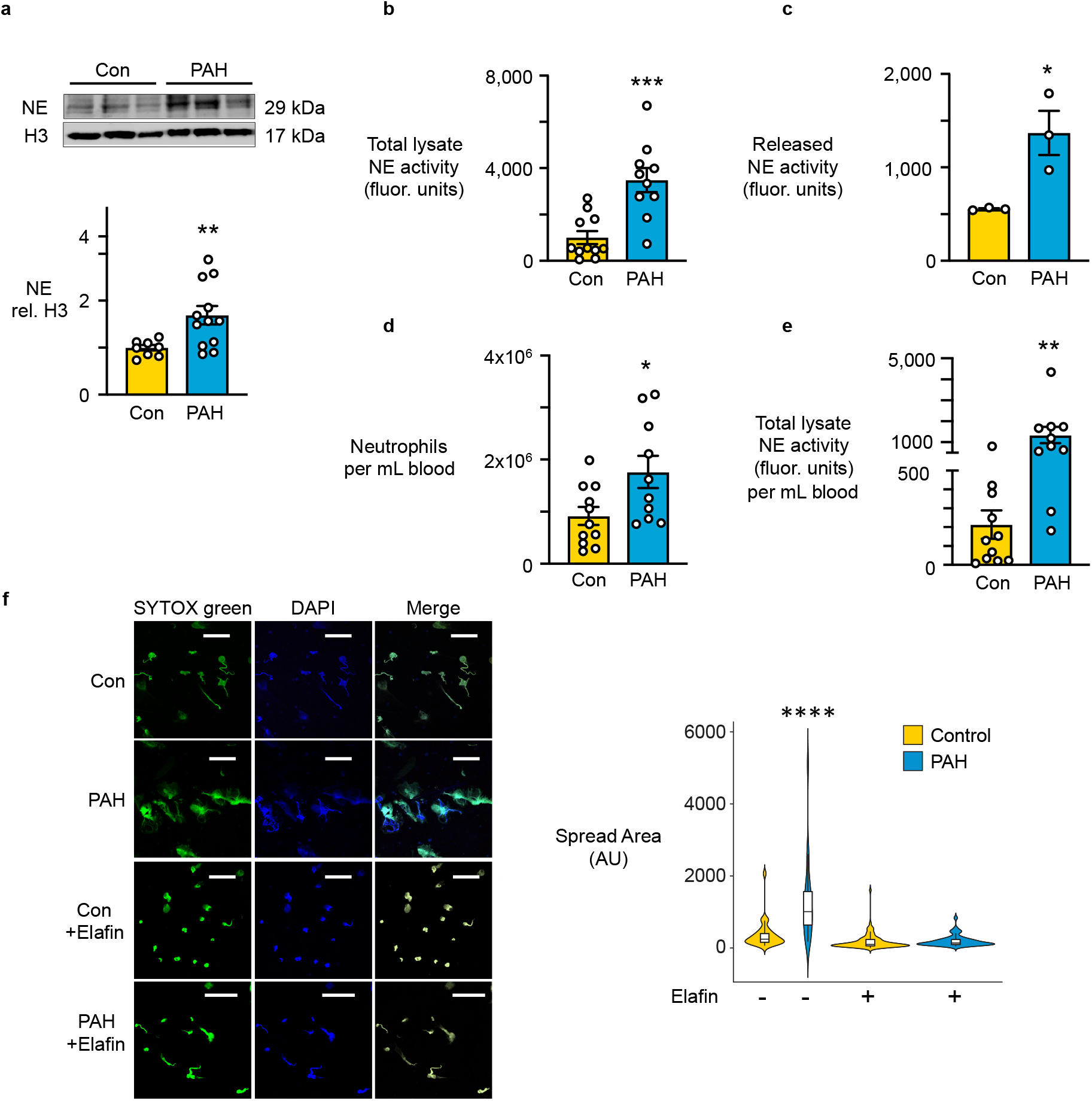
Increased NE protein and NE-mediated NETosis in PAH neutrophils. Neutrophils were isolated from peripheral blood of healthy donor controls (Con) and PAH patients (PAH) as described in the Methods. **(a)** Representative western immunoblot and quantification below, displaying NE relative to histone H3 (H3) in PAH vs. Con neutrophils (n=8 Con and n=12 PAH). **(b)** NE activity in neutrophil cell lysates as determined by production of BODIPY FL labeled fluorescent elastin fragments from self-quenching BODIPY FL-conjugated bovine neck ligament elastin (n=11 Con and n=10 PAH). **(c)** NE activity in the extracellular supernatant 2h after IL-8 stimulation of neutrophils (n=3). In a-e 5×10^6^ cells were used for each assay. **(d)** Number of neutrophils per mL blood in Con and PAH, determined using a Millipore scepter cell counter (n=11 Con and n=10 PAH). **(e)** Total NE activity calculated per 1mL blood. NE activity measured in 1b, relative to neutrophils counted per 1d, to yield NE per mL of blood. **(f)** Neutrophil extracellular traps (NET) were visualized and quantified by SYTOX green fluorescence, 60 min following PMA stimulation and treatment with vehicle (Veh) or the NE inhibitor elafin (1 µg/mL). Scale bar = 40µm. Violin plots represents variable distribution of 4 representative biological replicates (n = 24-104 cells). In a-e: Bars represent mean ± SEM. *p<0.05, **p<0.01, ***p<0.001 by unpaired Student t-test. In f for the violin plots a two-way ANOVA (normal distribution standard deviation) followed by Dunnets post hoc test ****p<0.0001.

Elevated NE activity was normalized to neutrophil counts that were also increased in PAH patients compared to control subjects (**Figure 1d**). Taking this into account, higher overall NE activity per milliliter of blood was present in PAH patients vs. healthy controls (**Figure 1e**). NE contributes to the release of neutrophil extracellular traps (NETs), by translocating from granules to the nucleus to cleave histones, causing chromatin decondensation and release^14^. NETosis incurs tissue damage, including endothelial cell (EC) injury^18^, through the exteriorization of chromatin accompanied by NE. Indeed, using SYTOX green staining of the spread area of exteriorized DNA, we showed heightened release of NETs upon PMA stimulation of PAH vs. control neutrophils that was inhibited by the NE inhibitor elafin^10^ (**Figure 1f**).

### PAH neutrophils exhibit increased adhesion and reduced migration

Increased NE production and NETosis suggested that other neutrophil functions could be altered or exaggerated in response to inflammatory stimuli. The functions considered were adhesion, migration and transendothelial migration. We observed heightened adhesion to a fibronectin substrate in PAH vs. control neutrophils (**Figure 2a**). Chemokinesis assessed by confocal microscopic live cell imaging of total distance moved was reduced in IL-8 stimulated PAH vs. control neutrophils (**Figure 2b, Supplementary Video 1**) as was chemotaxis, judged by longitudinal distance across the fibronectin substrate (**Figure 2c**). Similar to IL-8, stimulation with fMLP resulted in decreased chemotaxis in PAH vs. control neutrophils (**Supplementary Figure 2**), suggesting a process of impaired migration independent of a specific receptor activated pathway. We next investigated IL-8 stimulated neutrophil chemotaxis of PAH vs. control neutrophils across a monolayer of PAH or control pulmonary arterial endothelial cells (PAEC). While control neutrophils exhibited increased migration across PAH vs. control PAEC, PAH neutrophils displayed impaired migration across both control and PAH PAEC (**Figure 2d**).

**Figure 2.**
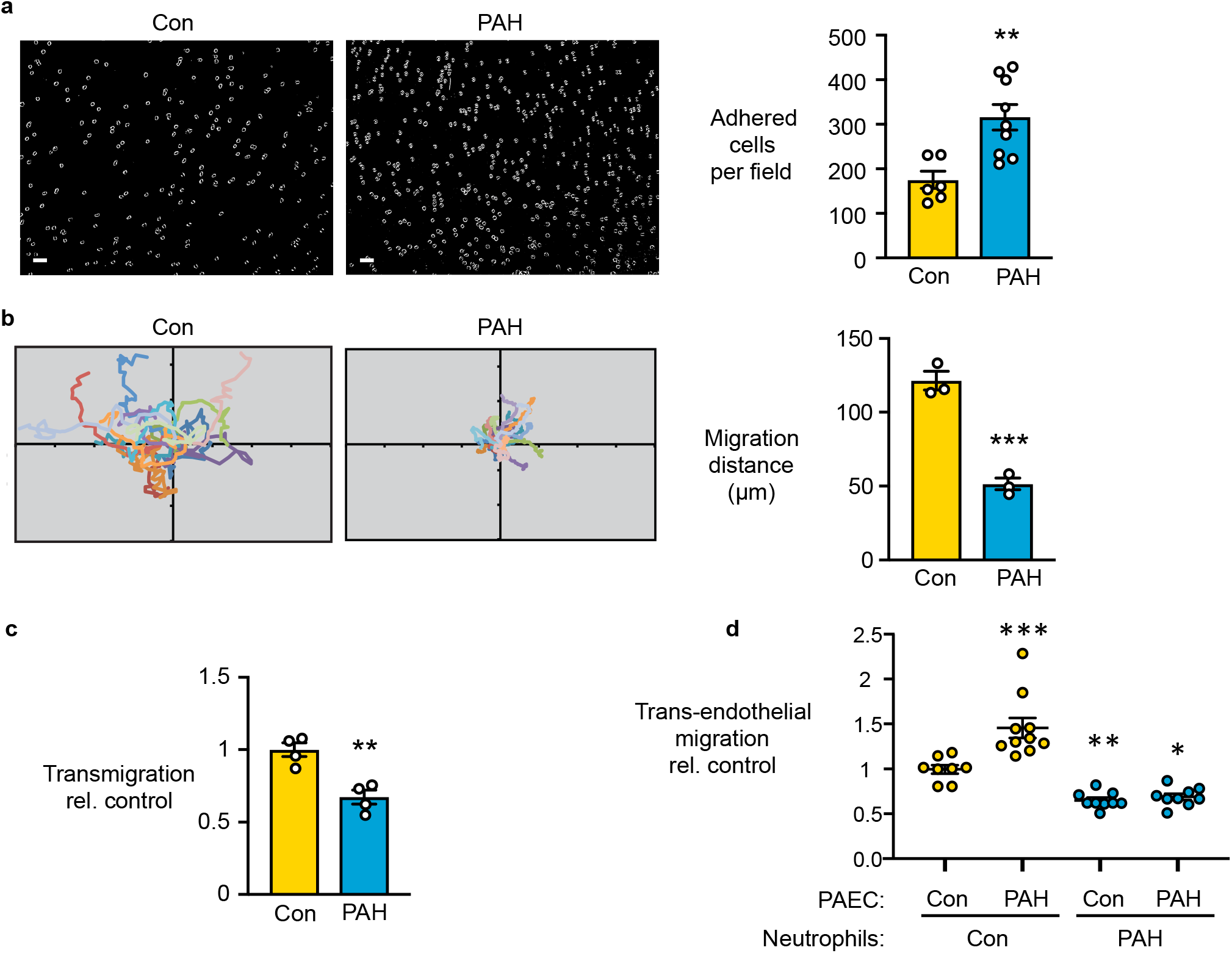
PAH neutrophils exhibit increased adhesion and reduced migration. (**a**) Neutrophils were incubated on fibronectin-coated coverslips for 5 min and the mean number of adherent cells was assessed from 3-5 randomly selected visual fields. Representative image of adherent neutrophils and quantification on the right (n=6 Con and n=9 PAH). Scale bar=40 µM. **(b)** Neutrophils were allowed to adhere to fibronectin coated coverslips for 5 min and then stimulated with 100 ng/mL IL-8 in RPMI for 30 min. Migration of PAH vs. Con neutrophils is illustrated as spider plots and distance migrated quantified (on the right) using MtrackJ (Image J). Quantification was performed for at least 10 migratory cells per field (n=3 fields). **(c)** Calcein-AM labeled PAH vs. Con neutrophils were plated on fibronectin coated transwell chambers, and then stimulated with 100 ng/mL IL-8. Transmigration was assessed after 60 min of stimulation (n=4). **(d)** Calcein-AM labeled PAH vs. Con neutrophils were incubated on Con or PAH PAEC monolayers grown to confluence in transwell chambers. Trans-endothelial migration of PAH vs. Con neutrophils across Con or PAH PAEC 60 min following IL-8 (100 ng/mL) stimulation (n=8-12). In a-c: Bars represent mean ± SEM. *p<0.05, **p<0.01, ***p<0.001 by unpaired Student t-test. In d: Range represent mean ± SEM by two-way ANOVA followed by Dunnets post hoc test. *p<0.05, **p<0.01, ***p<0.001.

### Proteomic analysis links increased vinculin to PAH neutrophil dysfunction

To investigate a mechanism that could explain the increase in NE activity and neutrophil adhesion and the decrease in migration, we carried out protein expression profiling by high-resolution mass spectrometry non-targeted proteomics. Principal component analysis (PCA) of the proteome revealed some overlap between PAH and control neutrophils (**Supplementary Figure 3a**), but 483 differentially regulated proteins were identified with a false discovery rate (FDR) of 10% (**Figure 3a**). Pathway Enrichment Analysis from IMPaLA, used to categorize functional changes, revealed proteins related to neutrophil degranulation, host interactions with HIV factors, adhesion, and transendothelial migration (**Figure 3b and Supplementary Figure 3b**). The increase in NE was evident by proteomic analysis with a p-value of 0.04, albeit an FDR of 11% (**Figure 3c**). Cathepsin G (CTSG), another serine protease present in azurophillic granules, was increased with a p-value of 0.006 and an FDR of 6% (**Figure 3c**). No significant differences were evident in other azurophilic proteases such as myeloperoxidase and azurocidin (**Supplementary Figure 3c**).

**Figure 3.**
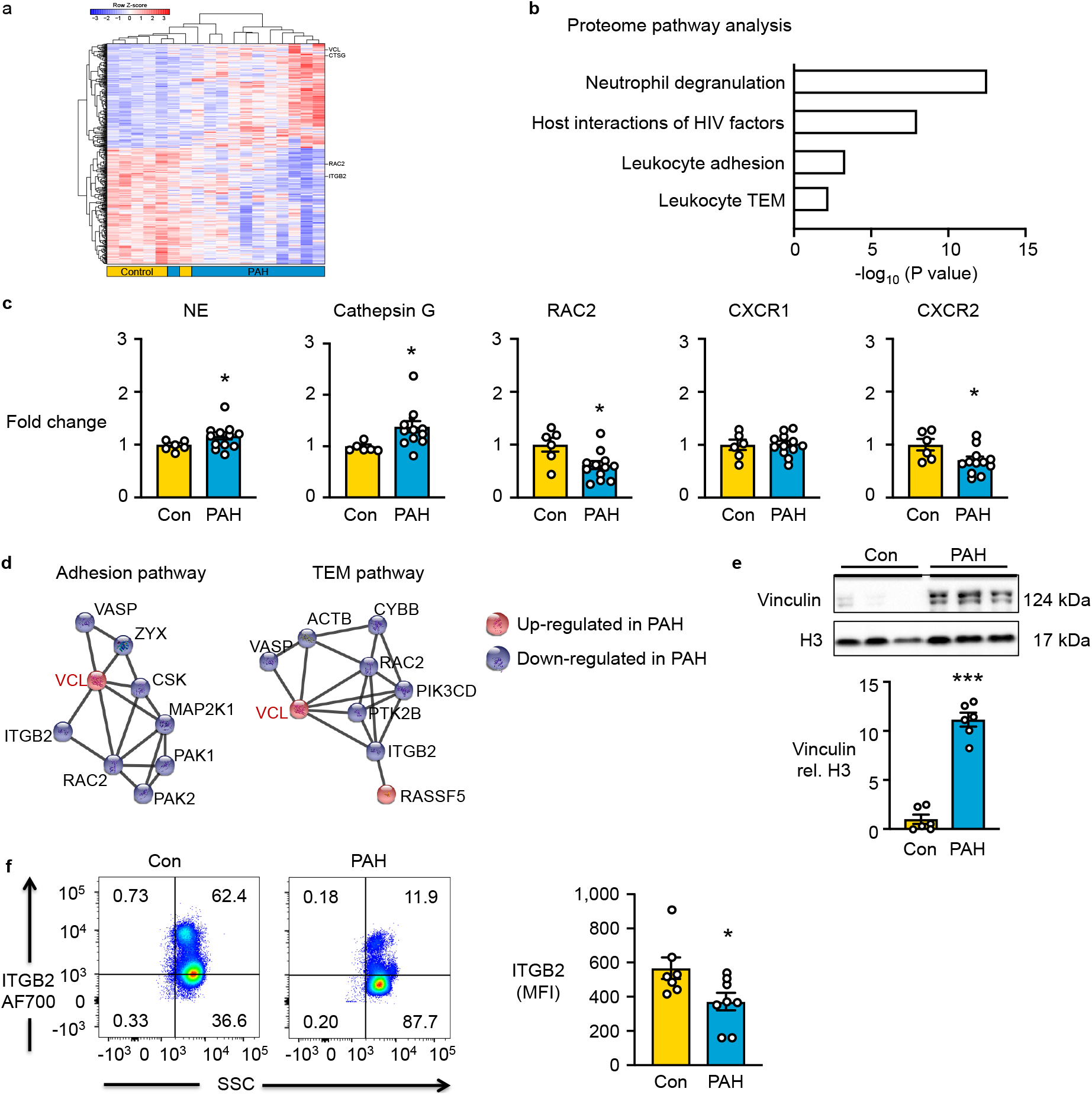
Proteomic analysis links increased vinculin to PAH neutrophil dysfunction. **(a)** Unbiased proteomic analysis of isolated PAH vs. Con neutrophils by mass spectroscopy. Heatmap depicts unsupervised hierarchical clustering of 483 differentially expressed proteins between the two groups (adjusted P value <0.10, n=6 Con and n=12 PAH. **(b)** Integrated Molecular Pathway-Level Analysis (IMPALA) reflecting significant differentially expressed proteins in PAH vs. Con neutrophils consistent with functional abnormalities identified in Figures 1 and 2. **(c)** Fold-change in NE, CTSG, RAC2, CXCR1, CXCR2 from proteomics analysis comparing PAH vs. Con neutrophils (n=6 Con and n=12 PAH). **(d)** STRING analysis of significant proteins in the Adhesion and TEM pathways, depicting protein up- or down-regulated in PAH vs. Con. **(e)** Representative western immunoblot and quantification below, validating the upregulation of VCL in PAH vs. Con neutrophils (n=6). **(f)** Validation and quantification of ITGB2 expression in neutrophils from Con or PAH by FACS analysis. Representative scatter graphs on the left, with quantification on the right (n=7 Con and n=8 PAH). In c, e, f: Bars represent mean ± SEM. *p<0.05, ***p<0.001by unpaired Student t-test.

Proteomic analysis also revealed reduced Ras-related Rho GTPase (RAC2) (p= 0.03, FDR, 9%) and C-X-C Motif Chemokine Receptor 2 (CXCR2) (p= 0.04, FDR 11%, **Figure 3c**) in PAH vs. control samples. RAC2 is an important mediator of neutrophil chemotaxis^22^, and IL-8 induces a migratory response through CXCR2 as reveiwed in ^23^. The reduction in RAC2 and CXCR2 could therefore contribute to decreased PAH neutrophil migration. CXCR1 is involved in neutrophil degranulation^24^ but was unchanged in the PAH vs. control samples. We next applied STRING analysis to find central protein nodes that could reveal key regulators in the adhesion and transendothelial migration pathways. Vinculin (VCL) presented as a central node in PAH samples that was linked to a reduction in integrin β2 (ITGB2), also known to play an integral role in adhesion and migration^25^ (**Figure 3d**). We confirmed the increase in VCL in PAH vs. control neutrophils by western immunoblot (**Figure 3e**) and the decrease in ITGB2 by Fluorescence Activated Cell Sorting (FACS) (**Figure 3f**). The increase in adhesion could be explained by heightened VCL, since VCL knockout neutrophils show reduced adhesion^26^. Moreover, fibronectin was used as the substrate for adhesion, and a strong VCL-fibronectin complex was previously reported^27^. The reduction in ITGB2 could explain reduced migration but not the heightened adhesion^28^.

### Transcriptomic analysis reveals an antiviral signature in PAH neutrophils

To elucidate mechanisms that could explain why NE and VCL are increased and ITGB2 is reduced in PAH vs. control neutrophils we conducted transcriptomic analyses using RNA-seq. The PCA analysis of the transcriptome revealed better separation of PAH and control neutrophils when compared with the proteomic analysis (**Supplementary Figure 4a**). Using an FDR below 5%, we identified 1,483 differentially expressed genes as displayed in the heat map (**Figure 4a**). By utilizing Pathway Enrichment Analysis from IMPALA, the top enriched pathway was related to neutrophil degranulation, consistent with the proteomic analysis. Other pathways were related to the immune system, innate immune system, and interferon signaling, suggesting an antiviral response (**Figure 4b and Supplementary Figure 4b**). As has been previously observed^29^, only a small subset of 34 genes and proteins were similarly up or down regulated on the proteomic and transciptomic analyses (**Supplementary Table 5**).

**Figure 4.**
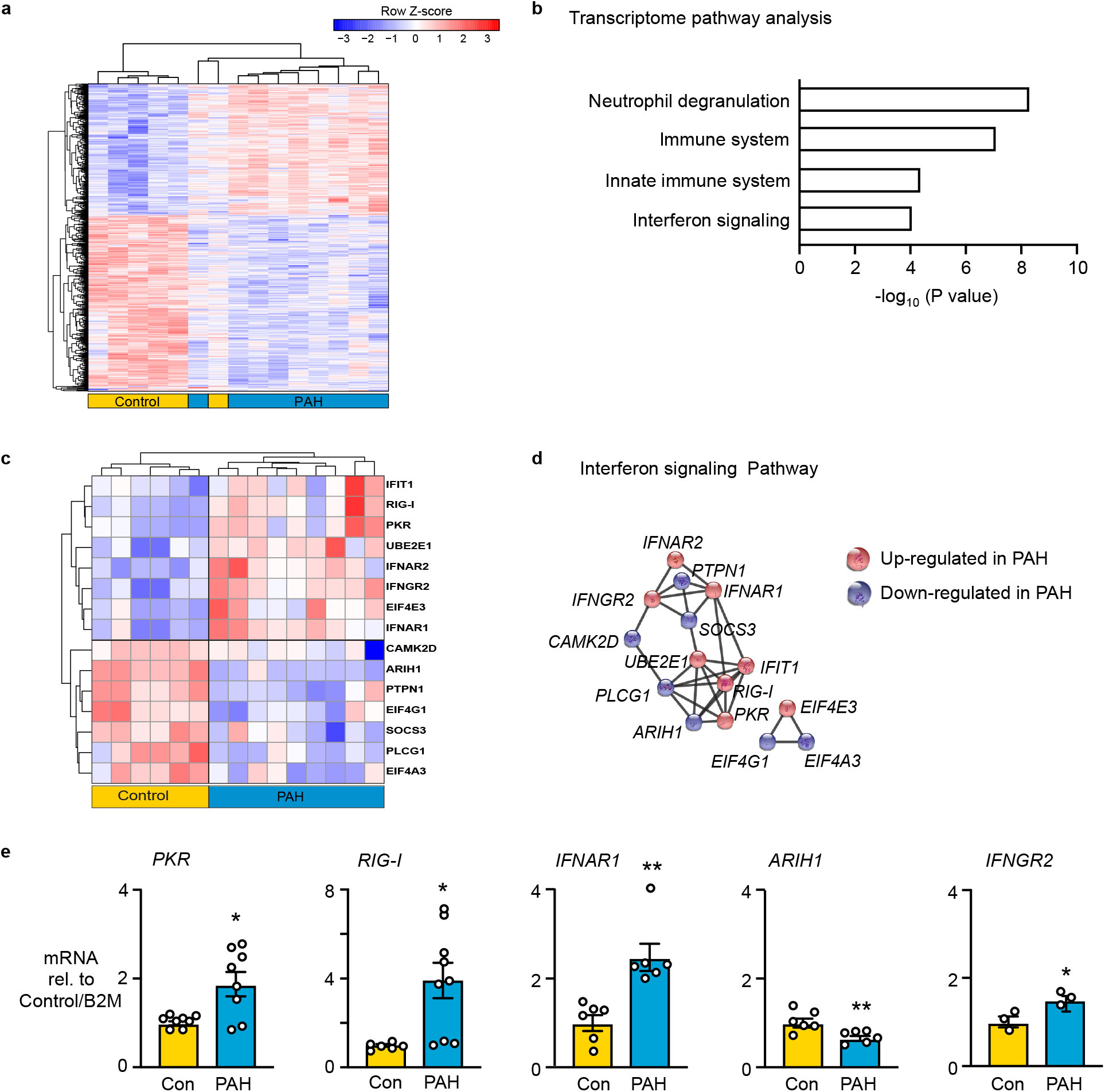
Transcriptomic analysis reveals an antiviral signature in PAH neutrophils. **(a)** Unbiased transcriptomics analysis was performed on neutrophils isolated from PAH vs. Con patients. Heatmap depicts unsupervised hierarchical clustering of 1,565 differentially expressed genes between the two groups (adjusted P value <0.05). (n=6 Con and n=9 PAH). **(b)** IMPaLA pathway analysis of significant genes in PAH vs. Con neutrophils. **(c)** Heatmap of significant differentially regulated genes in the interferon pathway, derived from the IMPaLA pathway analysis. **(d)** STRING analysis of significantly changed genes in the interferon pathway, depicting genes that are up- or down-regulated with key central nodes in PAH vs. Con. **(e)** mRNA expression by PCR, validating the changes in *PKR, RIG-I, IFNAR1, ARIH1*, and *IFNGR2 observed* in neutrophils isolated form Con or PAH patients. Bars represent mean ±SEM. *PKR* (n=8); *IFNR1* and *ARIH1* (n=6); *RIG-I* (n=6 Con and n=9 PAH); and *INFNGR*2 (n=3), *p<0.05, **p<0.01, ***p<0.001, by unpaired Student t-test.

There were 15 differentially expressed genes in the interferon signaling pathway as displayed in the heatmap (**Figure 4c**). STRING analysis was applied and the interferon-induced double-stranded RNA-dependent protein kinase *(PKR)* was determined to be a central node (**Figure 4d**). When activated, PKR limits viral replication during viral infection. An increase in *PKR*, retinoic acid-inducible gene I (*RIG-I)*, type I IFN receptor (*IFNAR1)*, and Interferon Gamma Receptor 2 (*IFNGR2)*, and a decrease in Ariadne RBR E3 Ubiquitin Protein Ligase 1 (*ARIH1)* are critical in mounting an effective antiviral immune response in neutrophils^30, 31^. We chose five interferon related genes for validation by qPCR and confirmed a PAH neutrophil vs. control increase in *PKR, RIG-I, IFNAR1*, and *IFNGR2* and a decrease in *ARIH1* (**Figure 4e**). Consistent with the antiviral signature in the transcriptome, VCL and ITGB2 are also implicated in an anti-viral response. The upregulation of VCL inhibits retrovirus infection in human cells^32^, and a reduction in ITGB2 in monocyte-derived macrophages results in impaired integrity of the human immunodeficiency virus (HIV)^33^.

### An increase in HERV-K envelope protein in PAH neutrophils is related to the antiviral signature and heightened NE

The anti-viral signature, evident from both the transcriptome and the proteome, and our previous studies in PAH monocytes and macrophages, led us to investigate whether an increase in endogenous retroviral elements might be present in PAH vs. controls neutrophils. We reported upregulation of HERV-K dUTPase in circulating monocytes of PAH patients and an increase in both the HERV-K dUTPase and the envelope protein in perivascular macrophages in PAH lung tissue sections^34^. As neutrophils are also of myeloid lineage, we assessed HERV-K proteins in PAH vs. control neutrophils and found an increase in the HERV-K envelope protein, but not in HERV-K dUTPase by western immunoblot analysis (**Figure 5a and Supplementary Figure 5a**). Consistent with the immunoblot, confocal analysis revealed heightened accumulation of HERV-K envelope protein in PAH vs. control neutrophils (**Figure 5b**).

**Figure 5.**
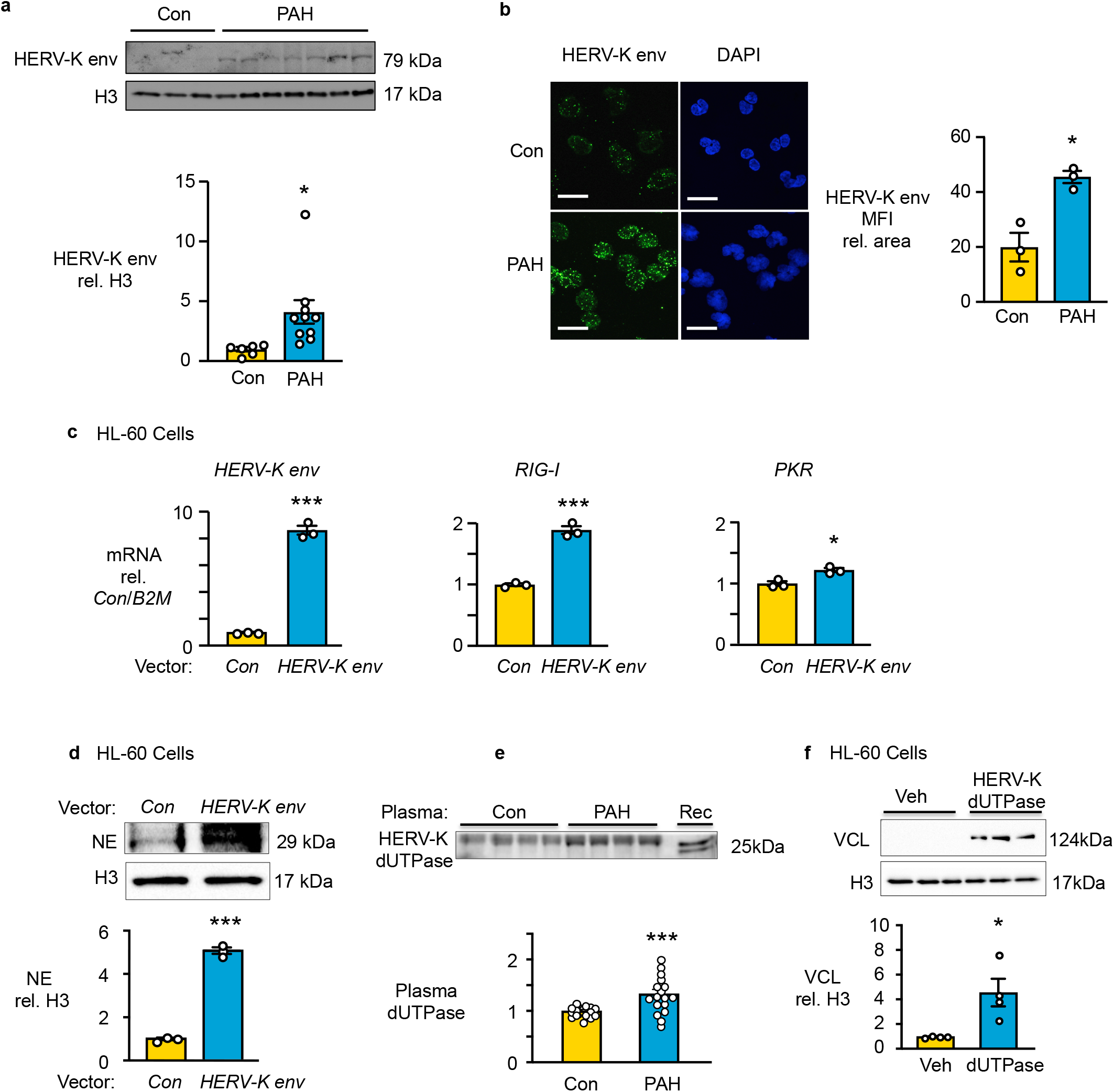
An increase in HERV-K envelope protein in PAH neutrophils is related to the antiviral signature and heightened NE. **(a)** Representative western immunoblot analysis and quantification of HERV-K envelope (HERV-K env) relative to histone H3 (H3) in PAH vs. Con neutrophils (n=6 Con and n=10 PAH). **(b)** Immunofluorescence microscopy visualization of HERV-K env protein (green) and nuclei by DAPI staining (blue) in Con and PAH neutrophils. Antibodies used for staining are described in the “Methods”. A range of z-stack images was collected for image analysis. Mean fluorescence intensity (MFI) and cell area were quantified via image j from 3-5 randomly selected visual fields (n=3). Scale bar = 20µm **(c-f)** HL-60 cells were cultured in RPMI complete media and transfected using HL-60 Cell Avalanche transfection reagent per manufacturer’s protocol. **(c)** RT-qPCR analysis of *HERV-K envelope, RIG-I*, and *PKR* mRNA in HL-60 cells overexpressing HERV-K env (n=3). **(d)** Representative western immunoblot and quantification of NE relative to H3 in HL-60 cells overexpressing HERV-K env (n=3). **(e)** Western immunoblot analysis and quantification of HERV-K dUTPase (dUTPase) in plasma collected from Con or PAH patients. Plasma was depleted of albumin, measured by BCA, and run on a gradient gel (n=17). (**f**) Representative western immunoblot analysis and quantification of VCL relative to H3 in dHL-60 cells, treated with 10 µg/mL HERV-K dUTPase or Veh (HERV-K dUTPase elution buffer) for 24 h (n=4). Bars represent mean ±SEM. *p<0.05 and ***p<0.001 by by unpaired Student t-test.

To determine whether the increase in HERV-K envelope was responsible for heighted NE and VCL as well as interferon signaling, HL-60 (a neutrophil promyelocytic cell line) was transfected with a HERV-K envelope vector or a control GFP vector, and gene expression was analyzed by qPCR (**Figure 5c**). We confirmed overexpression of *HERV-K envelope* that was directly related to upregulation of interferon genes *RIG-I* and *PKR* (**Figure 5c**) as well as increased NE protein expression (**Figure 5d**). There was, however, no significant elevation in VCL levels in HERV-K envelope vs. GFP transfected HL-60 cells (**Supplementary Figure 5b**).

As we detected increased circulating HERV-K dUTPase from PAH versus control plasma (**Figure 5e**), we evaluated whether HERV-K dUTPase produced by monocytes could be inducing VCL in PAH neutrophils in a cell non-autonomous manner. We added the recombinant form of HERV-K dUTPase to a neutrophil like cell line, differentiated HL-60 cells, and observed an increase in VCL (**Figure 5f and Supplementary Figure 5b**). There was, however, no significant upregulation of of interferon genes such as *PKR and RIG-1* or of NE protein, (**Supplementary Figure 5c and 5d**).

### CD66b^+^ PAH exosomes display increased NE and elevated HERV-K envelope

Transport of NE via neutrophil exosomes was recently shown to contribute to disease progression in a murine model of COPD^19^. We therefore used a CD66b antibody to purify neutrophil specific exosomes from PAH and control plasma after total exosome isolation using a nanofiltration-based exosome isolation tool, Exosome Total Isolation Chip (ExoTIC)^35^. Exosome size was confirmed by Nanosight (**Figure 6a**) and purity by transmission electron microscopy (**Figure 6b**). PAH exosomes exhibited an approximately 2-fold increase in NE protein, assessed by western immunoblot (**Figure 6c**) and by activity (**Figure 6d**) as well as a 4-fold increase in HERV-K envelope (**Figure 6c**).

**Figure 6.**
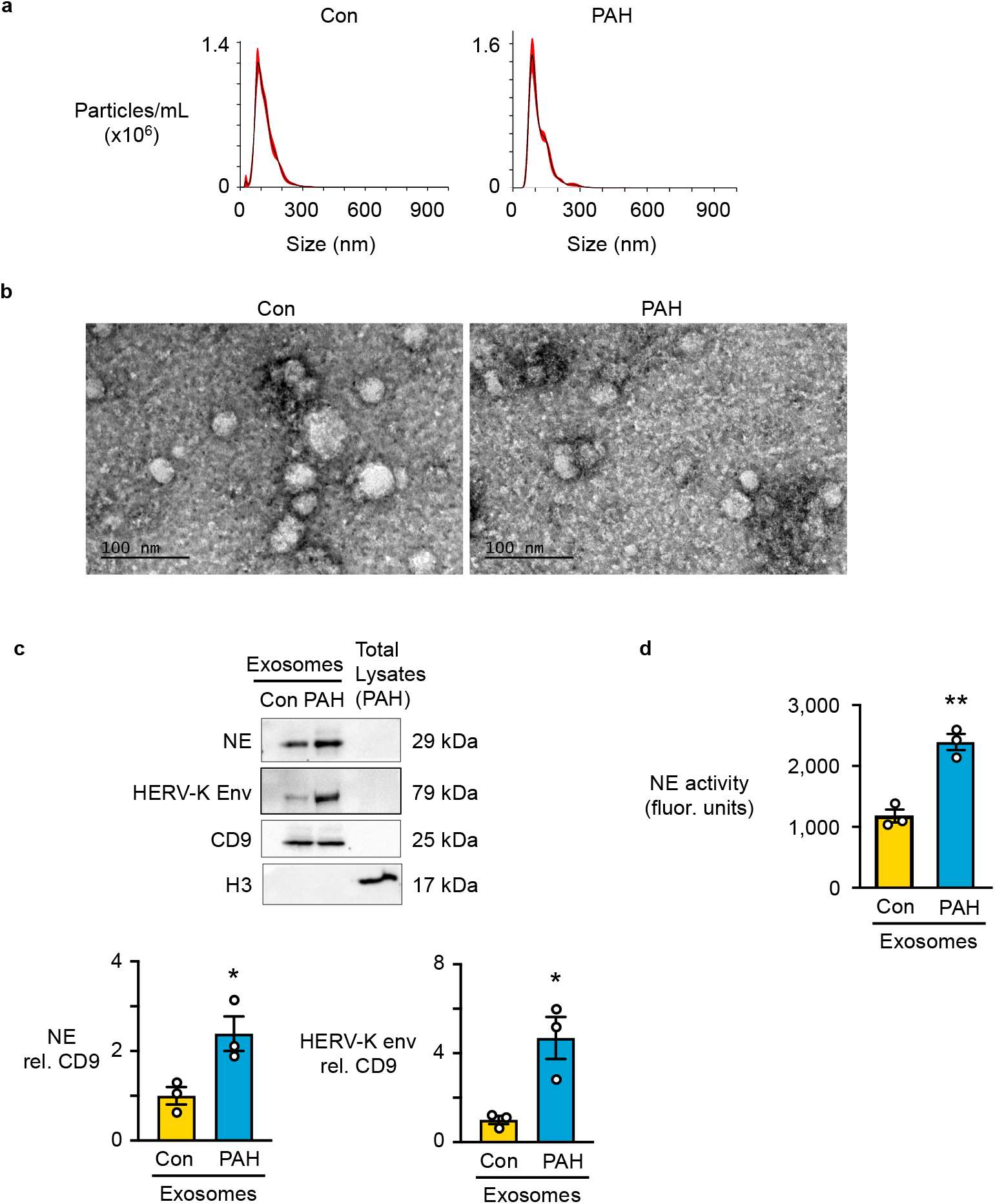
CD66b PAH exosomes display increased NE and elevated HERV-K envelope. Exosomes were isolated from plasma of 17 healthy donor controls and 17 PAH patients, using the ExoTIC device described under ‘Methods’. CD66b positive neutrophil exosomes were pulled down using anti-CD66b beads, from pooled exosomes of Con and PAH pooled plasma. **(a)** Size distribution of the pooled exosomes, determined using Nanosight. **(b)** Representative transmission electron microscopy (TEM) images of CD66b positive neutrophil exosomes derived from pooled plasma of PAH vs. Con patients. Scale bar=100 nm **(c)** Western immunoblot analysis and quantification of NE and HERV-K envelope from pooled PAH vs. Con neutrophil exosomes, relative to the exosome marker CD9. H3 from PAH neutrophil total lysate was used as a negative control. **(d)** NE activity in PAH vs. Con exosomes after 120 min incubation. NE was assessed by the production of BODIPY FL labeled fluorescent elastin fragments from self-quenching BODIPY FL-conjugated bovine neck ligament elastin. Bars represent mean ± SEM n=3 technical replicates of the pooled exosomes. *p<0.05, **p<0.01 by unpaired Student t-test.

### CD66b PAH exosomes cause pulmonary hypertension in a mouse model

We next determined whether pooled neutrophil exosomes isolated from PAH plasma could induce PAH in mice when compared to the same number of neutrophil exosomes isolated from control plasma. Neutrophil exosomes were given by tail vein injection, twice a week for five weeks to adult male mice (**Figure 7a**). This protocol is consistent with previous studies using exosomes from patients to model disease in mice^19^. Pulmonary hypertension was evident following injection of PAH but not control neutrophil exosomes, judged by decreased pulmonary artery acceleration time (PAAT) (**Figure 6b**), and increased right ventricular systolic pressure (RVSP), right ventricular hypertrophy (RVH), and peripheral pulmonary arterial muscularizaton (**Figure 7b-e**). Pretreatment of the exosomes with the NE inhibitor and antiviral agent elafin prevented the increase in RVSP and RVH (**Figure 7c and 7d**) as well as the increased muscularization of the pulmonary arteries (**Figure 7e**).

**Figure 7.**
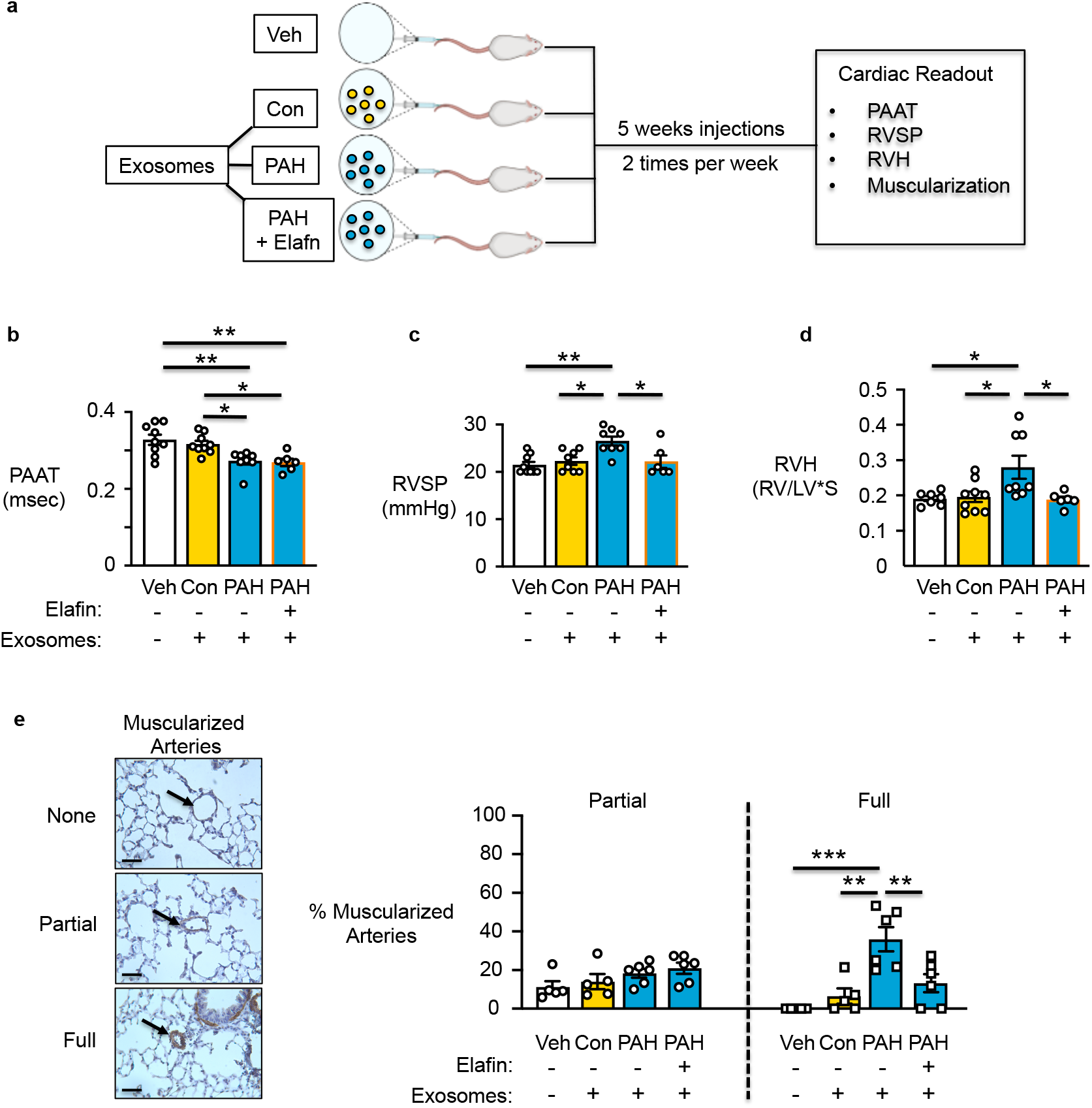
CD66b exosomes cause pulmonary hypertension in a mouse model. CD66b exosomes were isolated from plasma of PAH patients or healthy controls. Adult male mice (8 weeks) were injected with PAH vs. Con plasma exosomes (1.63 × 10^7^) at a volume of 100 µL. The PAH plasma exosomes were preincubated for 30 min with either elafin (0.02 mg/kg mouse weight in a volume of 4-6 µL from a 100 µg/mL stock solution), and or an equivalent volume of vehicle (PBS). The exosome suspensions were then injected twice a week, for 5 weeks. Hemodynamic function was evaluated 2 days after the last injection. **(a)** Illustration of animal protocol. **(b)** Pulmonary artery acceleration time (PAAT). **(c)** Right ventricular systolic pressure (RVSP). **(d)** Right ventricular hypertrophy (RVH). **(e)** Microscopy images of lung sections of the mice, labeled for αSMA (brown, smooth muscle cell marker) showing examples of nonmuscular, partially muscular and fully musularized vessels. Scale bar=40 µm. In b-e bars represent mean ± SEM. *p<0.05 **p<0.01, ***p<0.001 by one-way ANOVA followed by Dunnet’s post test comparing each group mean with the control group.

## DISCUSSION

Despite many advances in our understanding of the sequelae of chronic inflammation related to PAH pathology, the role of neutrophils has remained elusive. Here, we provide evidence of neutrophil dysfunction in PAH that we link to an antiviral response triggering innate immune defense mechanisms, that result in adverse remodeling of the pulmonary vasculature. We show that PAH neutrophils have increased NE and an associated propensity to form NETs. The increased adhesion and reduced migration that we also observed in PAH neutrophils can potentially amplify the damaging impact of NE release and NET formation. We relate the increased adhesion and reduced migration primarily to elevated levels of VCL despite reduced ITGB2^33^ and we attribute both features to an antiviral response^32, 33^ observed by transcriptomic analyses. Augmented retroviral HERV-K envelope was directly related to increased NE protein and activity and antiviral *PKR-RIG-1* mRNA levels. A non cell autonomous increase in HERV-K dUTPase, likely from monocytes, explained the increase in VCL. Elevation of NE and HERV-K envelope was evident in exosomes released from PAH neutrophils that induced features of pulmonary hypertension in mice. These observations reveal how an exaggerated innate immune response in neutrophils can contribute to the pathogenesis of PAH (summary schema, **Figure 8**).

**Figure 8.**
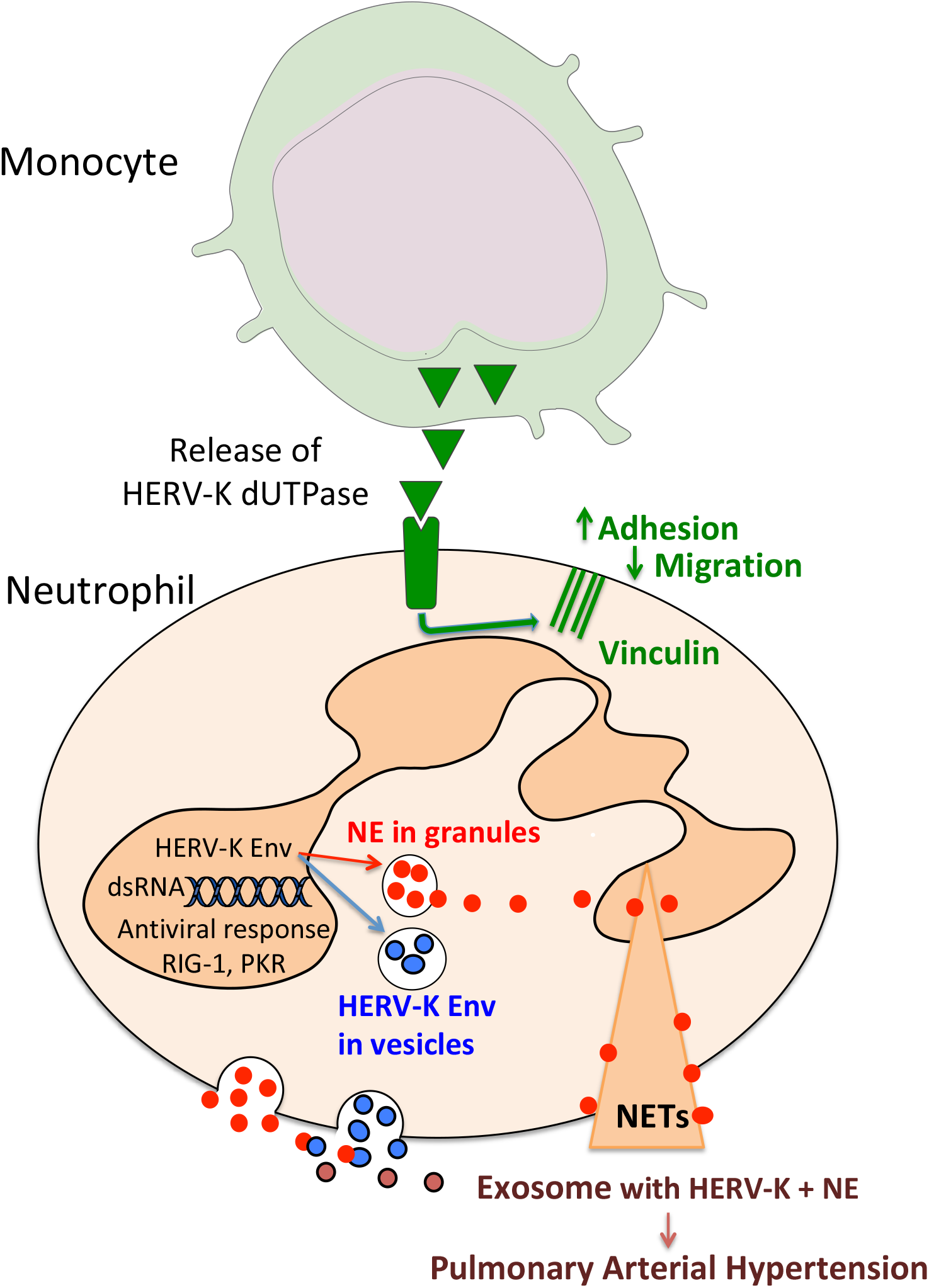
Schematic representation of PAH neutrophil. The secretion of HERV-K dUTPase from monocytes results in the upregulation of vinculin thereby increasing neutrophil adhesion and reducing migration. An increase in HERV-K envelope (Env) in the neutrophil is likely via double strated (ds) RNA required for the interferon response and heightened neutrophil elastase. The increase in neutrophil elastase promotes neutrophil extracellular traps NETs. Elastase released from granules associates with HERV-K env in exosomes causing pathologic features of pulmonary arterial hypertension.

Our previous studies attributed the increase of NE in PAH to its production by pulmonary arterial SMC in patients and experimental models^36^. In the present study, circulating neutrophils are increased approximately two-fold in PAH patients along with their greater than two-fold heightened expression of NE. This markedly increases the propensity for elastase mediated tissue damage, in part as a result of heightened predisposition to NET formation, previously described in PAH tissues^12^ and in part related to NE released in exosomes^19^.

Although it is known that NE resides on NETs and that NETs cause tissue damage^14^, we demonstrate a distinct relationship between the increase in NE and the propensity to form NETs in response to external danger signals phenocopied by PMA. The collateral damage of NETs in causing endothelial cell dysfunction^18^ has been related to NE^37^. While NE on NETS is shielded from natural circulating large molecule inhibitors such as alpha-1 antitrypsin^37^, our studies now provide evidence that small molecule inhibitors such as elafin can block NE activity and NET formation. NET markers (DNA, myeloperoxidase, and citrullinated histone H3) were previously identified In PAH lung tissue in the occlusive plexiform lesions, composed of inflammatory cells, aberrant endothelial cells and proliferative smooth muscle cells and is was shown that isolated neutrophil NETs triggered PAEC dysfunction^38, 39^.

Increased NE is generally associated with enhanced neutrophil adhesion as well as migration^16, 17^, as observed in a variety of lung diseases, including cystic fibrosis, severe asthma, chronic obstructive pulmonary disease and acute respiratory distress syndrome^40 41^. Although PAH neutrophils are characterized by increased adhesion, they exhibit reduced migration. This suggested that a non-NE mediated mechanism could be the cause of increased adhesion and reduced migration of PAH neutrophils. Proteomic analysis and western immunoblots confirmed an increase in VCL, a protein involved in the maturation of integrin-based focal adhesions, explaining both the reduced migration and increased adhesion. Consistent with our observations, previous studies reported that a reduction in neutrophil VCL resulted in decreased adhesion and increased migration^26^. Vinclulin suppresses ITGB2^42^, consistent with reduced migration as ITGB2 null mice exhibited reduced tissue emigration^28^. The reduction in CXCR2 and RAC2 might also contribute to the reduced migration^23^.

Surprisingly, the decrease in ITGB2 seems at odds with the increase in NE, since NE stabilizes adhesion of neutrophils by increasing the expression of ITGB2^16^, and NE is associated with both migration and adhesion of neutrophils, as reflected in studies in human neutrophils using a protease inhibitor or in NE-deficient mice^16, 17^. Thus, while the increase in VCL could explain enhanced adhesion and the reduction in ITGB2 could impact decreased migration in PAH neutrophils, these features appears to be NE independent. Another proteolytic enzyme that is increased in PAH neutrophils is cathepsin G. Although not elastolytic, an increase in cathepsin G can heighten the elastolytic activity of NE^43^ as shown in a model of aortic aneurysm^44^. In many lung diseases, such as COPD^19^, in addition to PAH^2, 10^, the breakdown of elastin by NE causes major tissue damage. The role of NE in adverse pulmonary vascular remodeling was previously reported in experimental pulmonary hypertension models where NE inhibition both prevented and reversed the disease (reviewed in ^1^).

A PAH neutrophil related anti-viral signature identified by transcriptomic analysis led to a mechanism that could explain increased NE and VCL consistent with an interferon innate immune response^45^. We investigated whether PAH neutrophils express high levels of endogenous retroviral RNA and protein of the HERV-K family, consistent with an observation we had made in PAH monocytes and macrophages^34^. HERVs are non-infectious remnants of ancient viral infections incorporated in our genome. They are highly expressed in embryonic stem cells as part of an innate immune response but undergo silencing in differentiated cells in response to extensive methylation of regulatory elements^46^. Expansion of retroviral RNA sequences with production of double stranded RNA and proteins can occur as part of an innate defense mechanism^47^ in response to systemic and environmental cues such as exogenous viruses, including human herpesviruses and HIV^48^. Although an increase in HERVs has been described in cancer^49^ and in autoimmunity and associated with an antiviral interferon response^45^, the mechanism for HERV upregulation in disease has been largely elusive. KAP1 is a methylase implicated in HERV methylation^50^ that is controlled by a BMP responsive lnc RNA, BORG^51^. So, it is possible that with reduced BMPR2 function, as occurs in PAH, BORG and KAP1 are reduced^51^. However, this does not explain why increased HERV-K is a feature of myeloid cells (monocytes and neutrophils) that we do not observe in vascular endothelial cells.

Overexpression of HERV-K envelope in a neutrophil cell line was sufficient to induce the production of NE. While the increase in NE likely represents the activation of an innate immune response, future studies will be of interest in delineating the molecular mechanism involved. In addition to NE, HERVs can mediate a chronic interferon response in association with the production of double stranded RNA^47, 52^, and interferon treatment is associated with the development of PAH^53^. Infection of monocyte-derived macrophages with HIV led to the upregulation of VCL, and as a consequence, VCL negatively affected the propagation of the virus^54^. Similar to HIV, recombinant HERV-K dUTPase also resulted in an increase in VCL, presumably by a similar mechanism.

Pathologic exosomes are recognized in many diseases, including autoimmunity^55^, cancer^58^, and COPD^19^. Recent studies support HERV-K delivery in tumor exosomes as a mechanism contributing to cancer^58^ and NE on exosomes as directly related to COPD^19, 56^. Here we identify an increase in both NE and HERV-K envelope in PAH neutrophil exosomes. In addition to these cell products, exosomes deliver multiple RNA transcripts and microRNAs, DNA, lipids, and proteins, hence their wide-range of effects in the initiation and perpetuation of disease, in their participation in the immune response, and in vascular dysfunction in association with tissue damage^57^. NE-rich exosomes, isolated from COPD patient bronchial lavage, caused degradation of airway structure and emphysema in a mouse, mimicking COPD. In this model, NE on exosomes was susceptible to inhibition by small molecules but evaded large protease inhibitors such as alpha-1 anti-trypsin^19^.

Studies in tumors demonstrate that HERV-K mRNA is packaged in exosomes that can be transferred to other cells^56^. In this study, recombinant elafin, a small 6 kDa protein, successfully blocked pulmonary hypertension induced by PAH exosomes. Elafin is a potent, naturally occurring NE inhibitor^10^, but it also has potent anti-inflammatory and antiviral properties that could explain its effect in preventing the adverse impact of HERV-K transferred in neutrophil PAH exosomes. For example, elafin is an NFκB inhibitor, and NFκB activation is associated with the interferon antiviral response^58^. Elafin also suppresses viral attachment and transcytosis^20^. Taken together, our studies reveal a mechanistic relationship between PAH and an increase in HERV-K mRNA and proteins that reprograms neutrophils by inducing an antiviral response. The production of large amounts of NE released in NETs locally and in exosomes can have long range adverse impact on vascular cells. These features can be targeted therapeutically and potentially reversed by Elafin.

## METHODS

### PAH patient and control samples

All human samples used were coded and all patients and controls signed an informed consent under protocols approved by the Institutional Review Board on Human Subjects in Medical Research at Stanford University. We included PAH patients between the ages of 23 and 80 and healthy controls between the ages of 27 and 60. Although patients were initially screened as having IPAH, 3/68 were re-classified as having drug and toxin (WHO Group 1 PAH)^21^ and 5/68 had comorbidities not responsible for PAH (**Supplementary Table 1**). Whole blood from PAH patients was obtained from the Stanford Pulmonary Arterial Hypertension biobank (IRB 14083), and blood from healthy volunteers was obtained from the Stanford biobank under the University sponsored precision health initiative (IRB 40869). Some experiments were carried out using de-identified blood from the healthy individuals, obtained from the Stanford Blood Center. PAEC used in the neutrophil trans-endothelial migration studies were from the Pulmonary Hypertension Breakthrough Initiative supported by NIH R24 HL123767 and the CMREF. Patients characteristics, demographics, and hemodynamics are shown in **Supplementary Tables 1 and 4**) and a list of the medications for each patient is decribed in **Supplementary Table 2**. Stanford biobank control samples used for the RNAseq and the untargeted proteomic LC/MS assay had echocardiographic studies performed to exclude cardiovascular disease (**Supplementary Table 3**).

### Reagents

See **Supplementary Table 6**.

### Neutrophil isolation

Human neutrophils were purified from peripheral blood using Miltenyi Biotec Macsxpress Neutrophil isolation kit, per manufacturer’s instructions. Purity of neutrophils after isolation was >95%, determined by cytospin preparation (Cytospin 4; Thermo Scientific, Waltham, MA) and Diff-Quick staining. Isolated Neutrophils were counted using a Scepter™ 2.0 Cell Counter (MilliporeSigma, Burlington, MA).

### HL-60 and dHL-60 cell culture

HL-60 cells were purchased from Sigma Aldrich and grown in complete media (RPMI 1640 medium supplemented with 5% FBS, 100 U/mL penicillin, and 100 μg/mL streptomycin) at 37°C in a 5% CO_2_ humidified environment. The cells used in the assays were between passages 10-30. To differentiate HL-60 cells into a neutrophil-like phenotype, 1.3% DMSO was added to the cell growth medium in which the cells were cultured for 6-7 days. On the final day, differentiated HL-60 cells (dHL-60) were centrifuged at 1,300 RPM for 5 min and resuspended in complete media. Viable cells were counted on a hemocytometer using trypan blue exclusion.

### HERV-K dUTPase protein purification

Recombinant HERV-K dUTPase protein was provided by Dr. Maria Ariza (Ohio State University). In some studies cells were incubated with recombinant HERV-K dUTPase at a concentration of 10 µg/mL for 24 hr prior to assay. The HERV-K gene encoding the dUTPase was cloned into the pTrcHis Topo TA expression vector and the sequence was verified by DNA sequencing analysis as previously described^59^. The purity of the expressed protein was assessed by SDS-PAGE and capillary-liquid chromatography nanospray tandem mass spectrometry performed at the Ohio State University Mass Spectrometry and Proteomics Facility. High-purity dUTPase preparations, free of contaminating DNA, RNA, lipopolysaccharide, and peptidoglycan, were used.

### PAEC cell culture

Primary small PAEC were harvested from the lungs of PAH patients undergoing lung transplantation, or from unused donor lungs (controls), obtained from the Pulmonary Hypertension Breakthrough Initiative (PHBI). Cells were cultured in EC medium supplemented with 5% FBS, 1% antibiotic/antimycotic, and 1% ECGS, and used between passages 4 and 7.

### NE activity

NE activity was measured using EnzChek Elastase Assay Kit (Thermofisher Scientific, Waltham, MA) according to the manufacturer’s instructions. For determination of intracellular NE activity, neutrophils were lysed with RIPA buffer followed by sonication. The solution was then centrifuged, and lysates were collected for NE activity analysis. For measurement of extracellular NE activity, neutrophils were stimulated with IL-8 (100 ng/mL) for 30 min at 37°C, and the supernatant collected for NE activity analysis.

### Neutrophil extracellular trap (NET) formation assay

Coverslips were pretreated with 100 µg/ml poly-d-lysine at 4°C overnight in phosphate buffered saline without calcium and magnesium (PBS-/-). Neutrophils were added to the coverslips for 5 min, and non-adherent cells washed away with PBS-/-. Phorbol myristate acetate (PMA) was added to the media at a final concentration of 100 nM for 60 min, with vehicle (PBS) or elafin (1 µg/mL) (Proteo-Biotech-AG, a kind gift of Dr. Oliver Wiedow, Kiel Germany). Cells were washed with PBS, and then stained with cell impermeable SYTOX green nucleic acid stain (3 µM, ThermoFisher Scientific) for 30 min. Cells were then washed and fixed with 4% PFA for 15 min at room temperature. Quantification of NETs was assessed by measuring the spread area of Sytox green positive DNA by capturing 3 images randomly from 4 biological replicates. Images were acquired using a Leica Sp8 (Leica, Wetzlar, Germany) confocal laser-scanning microscope with a 40X objective, and Leica Application Suite X software.

### Adhesion assay

Fibronectin (Sigma Aldrich, St. Louis, MO; 50 μg/mL) in PBS was applied onto 4 well chamber slides at 4°C, overnight. Purified neutrophils (1×10^5^) were added and incubated at 37°C for 5 min. The chamber slides were washed and at least 3 different fields of adherent cells in each chamber were imaged under light microscopy using a 10x objective and quantified.

### Transmigration assays

Transmigration assays were performed in a modified Boyden chamber using the Corning FluoroBlok 24-well, 3 µm pore size inserts. The inserts were coated with Fibronectin (Sigma Aldrich; 50 μg/mL) at 4°C overnight in PBS. The lower chamber was filled with DMEM supplemented with 1% BSA, 100 nM fMLP, or 100 ng/mL IL-8. Neutrophils (5×10^5^ cells/ml) were labeled with Calcein AM (3 µM) and plated on the fibronectin substratum. Neutrophils were allowed to migrate for 30 min in the presence of chemoattractant, and quantified via fluorescence plate reader at excitation and emission maxima of 494 nm and 517 nm, respectively.

### Migration (Chemokinesis)

The migration assay was used to determine real-time migration of neutrophils using time-lapse tracking. Coverslips were pretreated with 50 µg/mL fibronectin at 4°C overnight in PBS. Neutrophils were added to fibronectin pretreated coverslips for 5 min, non-adherent cells washed with PBS, and IL-8 added to the media at a concentration of 100 ng/mL. Cell migration was observed using a confocal laser-scanning microscope (FV1000, Olympus, Center Valley, PA) with a 40X objective. Frames were taken every 10 sec for 30 min and analyzed using ImageJ software.

### Transendothelial migration assay (TEM)

TEM was determined using the Corning FluoroBlok 24-well, 3 µm pore size inserts. PAECs were plated and grown to confluency on the inserts and washed with PBS. Neutrophils (5×10^5^ cells/mL) were labeled with Calcein AM (3 µM) and plated onto a pre-washed endothelial monolayer. IL8 (100 ng/mL) was dispensed into the lower chamber, and neutrophils were allowed to transmigrate through the endothelial monolayer towards the IL-8 chemoattractant. Neutrophil migration through the endothelial cell layer into the lower chambers was measured using a fluorescence plate reader at excitation and emission maxima of 494 nm and 517 nm, respectively.

### Flow cytometry

Neutrophils were incubated in blocking buffer (3% BSA in PBS) for 30 min at 4°C, then stained for surface expression using Alexa Fluor® 700 conjugated ITGB2 antibody for 1 h at 4°C. Neutrophils were washed with PBS supplemented with 1% BSA and fixed in a final concentration of 4% paraformaldehyde for 15 min at room temperature. The stained and fixed neutrophils were then suspended in PBS for data acquisition. Data was acquired using a Beckton Dickenson LSR II flow cytometer and analyzed using FlowJo version 10.4.2 software for mean fluorescent intensity (MFI). To ensure maximal purity, neutrophils were selected by using side scatter vs. forward scatter plots. Forward and side scatter gating strategy was used to determine cell size, granularity, and to exclude dead cells and debris from the neutrophil population. Unstained cells were used as a gating control for ITGB2 AF700.

### Immunofluorescence staining

Cells adherent to fibronectin were fixed with 4% paraformaldehyde for 15 min, permeabilized with 0.5% Triton X-100 for 1 min, and blocked in 5% FBS and 2% HSA in PBS-T (0.1%Tween 20) for 30 min. Cells were washed and then stained with HERV-K envelope primary antibodies (1:1000) (Ango) overnight at 4°C and Alexa Fluor 488 secondary antibodies (1:500) (Invitrogen) for 1 h at room temperature, washed, then mounted with DAPI Fluoromount-G (SouthernBiotech, Birmingham, AL). Confocal analysis was performed using a Leica Sp8 (Leica) confocal laser-scanning microscope with a 40X objective and Leica Application Suite X software.

### Reverse-Transcriptase qPCR (RT-qPCR)

Total neutrophil RNA was extracted using the Direct-zol RNA Kits (Zymo Research, Irvine, CA). The quantity and quality of RNA was determined using a a BioTek Synergy H1 Hybrid Reader (Winooski, VT). qPCR was performed using 5 μL Powerup SYBR green PCR Master Mix (Applied Biosystems, Foster City, CA), 2 μL of dH_2_O and 2 μL of cDNA sample in a 10 μL reaction. Each measurement was carried out in duplicate using a CFX384 Real-Time System (Bio-Rad, Hercules, CA). The PCR conditions were: 50°C for 2 min, 95°C for 10 min, followed by 44 cycles of 95°C for 15 sec, and 60°C for 60 sec. Primer sequences used are listed in **Supplementary Table 7**. Gene expression levels were normalized to beta-2-microglobulin (B2M).

### Transmission Electron Microscopy

Pooled exosome samples were prepared for electron microscopy in PBS. Grids were glow discharged for 20 sec in a Denton Desktop Turbo III vacuum system and then covered with 5 µL of sample and allowed to adsorb to a 300 mesh copper grid with formvar and carbon coating (Cat# FCF300-CU) for 3 min. The sample was washed by touching the grid sample side down with two drops of water. Uranyl Acetate (1% in water) was then dripped through a 0.2 µm mesh syringe filter over the grid, allowing the third drop to remain on the grid for 1 min, and most of the stain to wash away. Grids were observed using a JEOL JEM-1400 120kV electron microscope, and images acquired using a Gatan Orius 2k x 4k digital camera.

### Western Immunoblot

Neutrophils were isolated and washed three times with ice-cold PBS, and neutrophil lysates prepared by adding lysis buffer (10 mM Tris-HCI and 1% SDS), then boiled for 10 min before centrifugation. Protein concentration was determined by the BCA assay (Thermo). Equal amounts of protein were separated by SDS-PAGE and transferred onto nitrocellulose membranes. Membranes were incubated overnight at 4° C with antibodies against vinculin (1:1000), NE (1:500), HERV-K envelope (1:1000), H3 (1:3000), HERV-K dUTPase (1:500) or CD9 (1:1000), in 5% BSA-PBS containing 0.1% Tween-20. Secondary antibodies were added according to manufacturer’s protocol and membranes were imaged using chemiluminescence reagent Clarity Western ECL Substrate (BioRad laboratories, Hercules, CA) and a BioRad ChemiDoc XRS system. Densitometric quantifications were performed using Image Lab software version 5.2.1.

### Plasmid transfection

HL-60 cells were transfected with HERV-K envelope vector (VectorBuilder, Chicago, IL) or EGFP control vector (VectorBuilder) using HL-60 Cell Avalanche™ Transfection Reagent per manufactures instructions (EZ biosystems, College Park, MD).

### Exosome isolation from plasma and CD66b pulldown

Human plasma (500 µL) was prepared from blood of 17 separate healthy donor controls and 17 PAH patients. Exosomes were isolated from the plasma using the Exosome Total Isolation Chip (ExoTIC)as described in ^35^. ExoTIC-harvested exosomes were further purified using CD66b/CEACAM 8 antibody (Sigma), a specific human neutrophil marker, which was conjugated to beads using the Dynabeads Antibody Coupling Kit per manufacture instructions (ThermoFisher). Exosomes were co-incubated with magnetic beads conjugated with a CD66b/CEACAM 8 antibody in a shaker for 18 h at 4°C. Using a magnet, unbound exosome supernatant was removed and captured beads were washed. To dissociate the beads from the CD66b/CEACAM 8 positive exosomes, an acid wash was performed by adding 200 µL of 50 mM pH 2.5 glycine solution to tube containing beads for 10 min, followed by addition of 70 µL 1M pH 7.5 Tris buffer to neutralize the solution. The purity, size, and concentration of the eluted exosomes were evaluated by nanotracking analysis on a Nanosight NS300 (Malvern, State). Exosomes from the 17 healthy donor controls were pooled, and the exosomes from the 17 PAH patients were pooled, and equal concentrations of the pooled exosomes were aliquoted and stored in −80°C for further experiments.

### Untargeted Proteomics from Neutrophils by LC-MS

#### Sample preparation

Proteomics was evaluated using neutrophils isolated from 6 healthy control subjects and 12 PAH patients. Human neutrophils (5×10^6^) were lysed in 10 mM Tris-HCI and 1% SDS, and 40 µg of protein loaded and concentrated by SDS-PAGE at 100 volts for 10 min. After Coomassie blue staining, the protein bands from each sample were excised with a clean razor blade. Proteins were reduced, alkylated and digested overnight using Trypsin/Lys-C and peptides were extracted from the gel. Peptide extracts were then dried in a speed vac, reconstituted in 100 mM TEAB and protein concentrations were measured using the BCA method. Peptides were labeled with TMT 10plex reagent (Thermo Fisher) as instructed by the vendor and subsequently combined at equal amount. Each pool contained 9 samples and one reference pool sample.

#### Data acquisition

Tryptic peptides were separated in 2 dimensions on a MClass 2DnLC (Waters, city, state). First dimension consisted of a reverse phase chromatography at high pH, followed by an orthogonal separation at low pH in the second dimension. In the first dimension, the mobile phases were 20 mM ammonium formate at pH 10 (Solvent A) and 100% acetonitrile (Solvent B). Peptides were separated on a Xbridge 300 µm x 5 cm C18 5.0 µm column (Waters) using 15 discontinuous step gradients at 2 µL/min. In the second dimension, peptides were loaded to an in-house packed 100 µm ID/15 µm tip ID x 28 cm C18-AQ 1.8 µm resin column with buffer A (0.1% formic acid in water). Peptides were eluted using a 180-min gradient from 5% to 40% buffer B (0.1% formic acid in acetonitrile) at a flow rate of 600 nL/min. The LC system was directly coupled in-line with an Orbitrap Fusion Lumos Mass Spectrometer. The source was operated at 1.8-2.2 kV to optimize the nanospray with the ion transfer tube at 275°C. The mass spectrometer was run in a data dependent mode. Full MS scan was acquired in the Orbitrap mass analyzer from 400-1500 m/z with resolution of 120,000. Precursors were isolated with an isolation window of 0.7 m/z and fragmented using CID at 35% energy in ion trap in rapid mode. AGC target was 10^4^ and the maximum injection time was 100 ms. Subsequently, 8 fragment ions were selected for MS3 analysis, isolated with an m/z window of 1.6, and fragmented with HCD at 65% energy. Resulting fragments were detected in the Orbitrap at 60,000 resolutions, with a maximum injection time of 150 ms or until the AGC target value of 10^5^ was reached.

#### Data processing

Raw data were processed using Proteome Discoverer software PD2.1 (Thermo) with precursor and fragment ion mass tolerance of 10 ppm and 0.6 Dalton, respectively, for database search. The search included cysteine carbamidomethylation as a fixed modification. Acetylation at protein N-terminus, methionine oxidation and TMT at peptide N-terminus and lysine were used as variable modifications. Up to two missed cleavages were allowed for trypsin digestion. Only unique peptides with a minimum of six amino acids were considered for protein identification. The peptide false discovery rate (FDR) was set at 1%. Data was searched against the uniprot human proteome database. Quantitative results at the protein level were reported relative to the reference pool sample. Batch effect was corrected by applying median-normalization and proteins detected in less than 2/3 of the samples were discarded. Differential analysis was performed on log_2_-transformed data using a two-sided Welsh t test. Proteins were considered significant with a q-value below 0.10. Functional correlation networks were plotted using the webtool STRING (http://string-db.org).

### RNA-seq analysis pipeline

#### Sample Preparation

RNA sequencing (RNA-seq) was evaluated using neutrophils from 6 healthy control subjects and 9 PAH patients. Total RNA was prepared for sequencing using the Takara Bio SMARTer: SMARTer Stranded Total RNA-Seq Kit v2 - Pico Input Mammalian kit according to the manufacturer’s instructions.

#### Data acquisition and processing

Pooled libraries were sequenced using the HiSeq 2500 sequencer (Illumina, San Diego, CA, SA) at the Stanford Center for Genomics and Personalized Medicine facility. Healthy donor controls and PAH samples were sequenced in each pooled lane to correct for potential batch effect. Sequencing data were demultiplexed and converted into fastq files using Illumina’s bcl2fastq conversion software (v2.20). Quality and adapter trimming along with filtering of rRNA reads was performed using BBDuk (v37.22). cDNA sequences of protein coding and lncRNA genes were obtained from human genome assembly GRCh38 (ens85). Paired-end 100bp reads were mapped to the indexed reference transcript using kallisto 0.43.0 and the –fr-stranded option^60^. DEseq2 normalized fold-changes were used to estimate differential gene expression between control and PAH neutrophils using the ‘DESeq2’ R package (DESeq2 v1.16.1)^61^. We identified 1565 significantly differentially expressed genes at FDR <5% (896 downregulated, 669 upregulated). The heatmap of expression across samples for significant genes was plotted using the R package ‘pheatmap’ v1.0.8.

### Mouse model for the induction of pulmonary hypertension by neutrophil exosomes

The Animal Care Committee at Stanford University approved all experimental protocols used in this study, following the published guidelines of the National Institutes of Health and the American Physiological Society. Isoflurane anesthesia (1.5%, 1 L/min oxygen) was used during these procedures. Adult male mice (8 weeks of age) were given tail-vein injections twice weekly for five weeks of previously aliquoted and frozen exosomes (1.63×10^7^) from pooled healthy donor control or PAH plasma brought to room temperature. Male mice were used for comparison wih previous studies from our group ^34^. Prior to injection, the PAH exosomes were incubated for 30 min with either elafin in PBS (0.02 mg/kg mouse weight in a volume of 4-6 µL from a 100 µg/mL stock solution) or an eqivalent volume of PBS. Thereafter, cardiac function, right ventricular systolic pressure, and right ventricular hypertrophy were assessed as previously described^34^. Cardiac function was measured by Vivid 7 ultrasound machine (GE Healthcare, Pittsburgh, PA) and 13-MHz linear array transducer. Isoflurane anesthesia (1.5%, 1 L/min oxygen) was used during these procedures for RVSP and it was measured by inserting a 1.4F Millar catheter (Millar Instruments, Houston, TX) via right jugular vein. Data were collected by Power Lab Data Acquisition system (AD Instruments, Colorado Springs, CO) and analyzed by LabChart software (AD Instruments, Colorado Springs, CO). After hemodynamic measurements, the lungs were flushed with saline. Right ventricular hypertrophy was assessed by the weight ratio of the RV to LV plus septum. Right lungs were snap-frozen in liquid nitrogen and stored at −80° C. Left lungs were fixed with 10% formalin for histology. Paraffin embedded and formaldehyde fixed lung section were stained for alpha-SMA (Sigma-Aldrich) using the Dako Animal Research Kit (Dako, Denmark) per manufactures instructions as previously described. Light microscopic images were acquired for quantification of percent partially and fully muscularized peripheral alveolar duct and wall. Three fields (200x magnification) were assessed (one in each of three lung sections/mouse) as previously described^34^. Quantification of muscularization was conducted in a blinded manner.

### Statistical Analysis

Individual data points are shown, with bars representing mean±standard error of the mean (SEM). Statistical significance was determined by one-way or two-way ANOVA followed by Dunnets multiple camparison test, or two-sided unpaired t-test analysis as indicated in the figure legends. Statistical analysis was performed using GraphPad Prism (v8.4.1) or R package ‘impute’ (v1.52.0). Pathway enrichment analysis was conducted using the webtool IMPaLA (https://impala.molgen.mpg.de/).

## ACKNOWLEDGMENTS

We thank Drs. Angela Rogers and David Cornfield at Stanford University for their critical input of our work; Devon Kelley, Matthew A. Bill, Jordan Burgess, Audrey Inglis, Alex Aaron Yacob, and Divya Rajmohan for collecting PAH patient blood samples, and Kalyani Anil Boralkar and Shadi Peighambari Bagherzadeh for the collection of healthy donor control blood samples. We greatly appreciate Dr. Michal Roof for her input, editorial, and technical assistance. We thank Ms. Ruiqi Jian for proteomic sample preparation, Dr. Tushar Desai for the use of the Leica confocal microscope, Dr. Dan Bernstein for the use of his cardiac phenotyping equipment, and Ms. Michelle Fox for her administrative help.

This work used the Genome Sequencing Service Center of the Stanford Center for Genomics and Personalized Medicine Sequencing Center, supported by NIH grants S10 OD025212 and P30 DK116074 (MPS). Proteomic peptide isolation was performed under the guidance of Anna Okuma, Ryan Leib, Christopher M. Adams, and Roasa Mehmood at the Vincent Coates Foundation Mass Spectrometry Laboratory, Stanford University. The mice histology sectioning was performed by the Department of Comparative Medicine Animal Histology Services (AHS). The transmission electron microscopy imaging was performed by the Cell Sciences Imaging Facility (CSIF), which is partly supported by ARRA/NCRR grant S10 RR026780. Flow samples were run at the Stanford Shared FACS Facility. The Pulmonary Hypertension Breakthrough Initiative (PHBI), funded by NIH/NHLBI R24 HL123767 and the Cardiovascular Medical Research and Education Fund (CMREF) UL1RR024986, provided PAEC from PAHG patients and unused donor lungs.

## Funding Sources

NIH-NHLBI grants P01 HL108797 (MR), R01 HL122887 (MR, MRN and MPS) R01 HL074186 (MR), NIH-NIAID grant R01 A1084898 (MEA), NIH-NHLBI grant R01 HL 138473 (MR and MRN, and R01 HL074186 (MR). ST was supported by T32 Fellowship in Pulmonary Medicine T32 HL129970-02 (PI: MRN) and an NIH/NHLBI Research Diversity Supplement P01 HL108797-04W1. SI was supported by AHA award 20POST35080009; SO was supported by a grant from Mie University Graduate School of Medicine; JRM by the California TRDRP of the University of California award 27FT-0039, and by the Netherlands Heart Foundation award 2013T116; DPM is supported by NIH-NHLBI grant K99 HL 1450970; MG by NIH/NHLBI grant K99/R00 HL135258; KM by fellowships from Japan Heart Foundation/Bayer Yakuhin Research Grant Abroad and The Uehara Memorial Foundation. AART was supported by a British Heart Foundation-Fulbright scholar award and Intermediate Clinical Fellowship (FS/18/13/33281). BAB was supported by NIA R00 AG049934. AJS was supported by NIH/NHLBI grant K23 HL151892. MR is also supported in part by the Dunlevie Chair in Pediatric Cardiology at Stanford University.

## CONTRIBUTIONS

ST and MR conceptualized the project. ST performed experiments, analyzed data, edited the figures, and wrote the manuscript. KC, BB, and ST performed omics data processing and analysis. LJ performed omics data acquisition. ST, LW, SI, SO, TS, and JK performed mouse cardiac hemodynamics. FH, RTZ, DH, PDR, and AJS provided human cardiac hemodynamics and samples. UD supervised exosome isolation, MO performed exosomes isolation, and MO and DM performed Nanosight analysis. SM and ST performed tail vein injections for animal experiments. MG, JM, AC, AART, KM assisted in experiments regarding cell adhesion (MG), confocal imaging (JM), neutrophil isolation (AC), co-cultures (KM), and transfections (AART) MEA provided purified HERV-K dUTPase protein and anti-HERV-K dUTPase Ab. MPS and MRN assisted in the supervision of the project. MR supervised the project and helped prepare the manuscript. All authors discussed the results and commented on the manuscript.

## COMPETING INTERESTS

None

## DATA AVAILABILITY

RNAseq data will be deposited into database of Genotypes and Phenotypes (dbGaP) (in Progress). Proteomic will be deposited to the ProteomeXchange Consortium via the PRIDE partner repository (in Progress).

## Code availability

Code related to RNA-seq data analysis is available on the Benayoun lab github.

## Notes

### Competing Interest Statement

The authors have declared no competing interest.

