## Supplemental Figures and Tables for "Endogenous Retroviral Elements Generate Pathologic Neutrophils and Elastase Rich Exosomes in Pulmonary Arterial Hypertension"

### SUPPLEMENTARY FIGURES AND LEGENDS

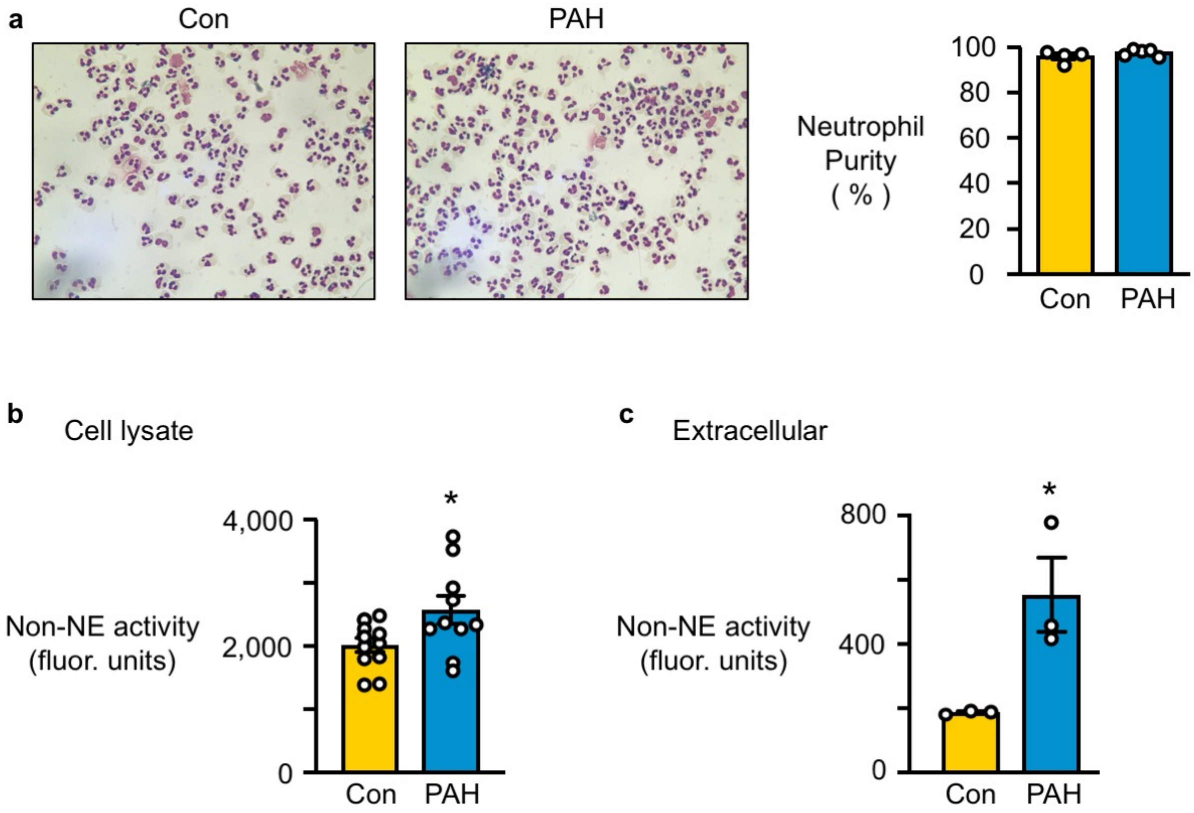


**Supplementary Figure 1. Non-NE activity and validation of neutrophil purity.**

**(a)** Neutrophils were isolated from circulating human blood, and the purity of the preparation was determined by microscopy following cytospin preparation (Cytospin 4; Thermo Scientific) and Diff-Quick staining (n=4 Con or 5 PAH). **(b)** Elastase activity that is not due to NE in neutrophil cellular lysates after 2 h. Non-NE activity was assessed by the level of BODIPY FL labeled fluorescent elastin fragments produced from self-quenching BODIPY FL-conjugated bovine neck ligament elastin (n=11 Con or n=10 PAH). **(c)** Non-NE activity was measured in extracellular supernatant of PAH and control neutrophils 2 h after IL-8 stimulation (n=3). Bars represent mean ± SEM. *p<0.05 by unpaired Student t-test.

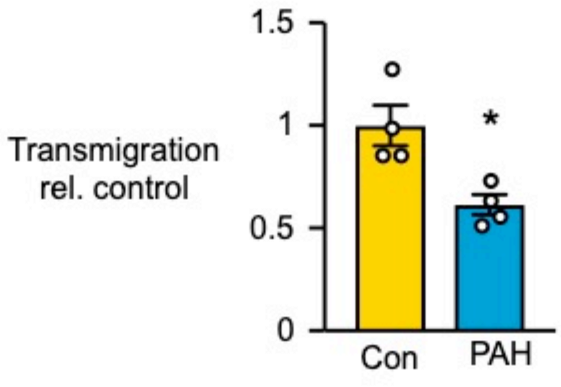


**Supplementary Figure 2. PAH neutrophils exhibit reduced migration**

Neutrophils were labeled with Calcein-AM, plated on fibronectin coated trans-well chambers, and stimulated with 100 nM f-MLP. Transmigration was assessed after 60 min of stimulation (n=4). Bars represent mean ± SEM. *$p<0.05$ by unpaired Student t-test.

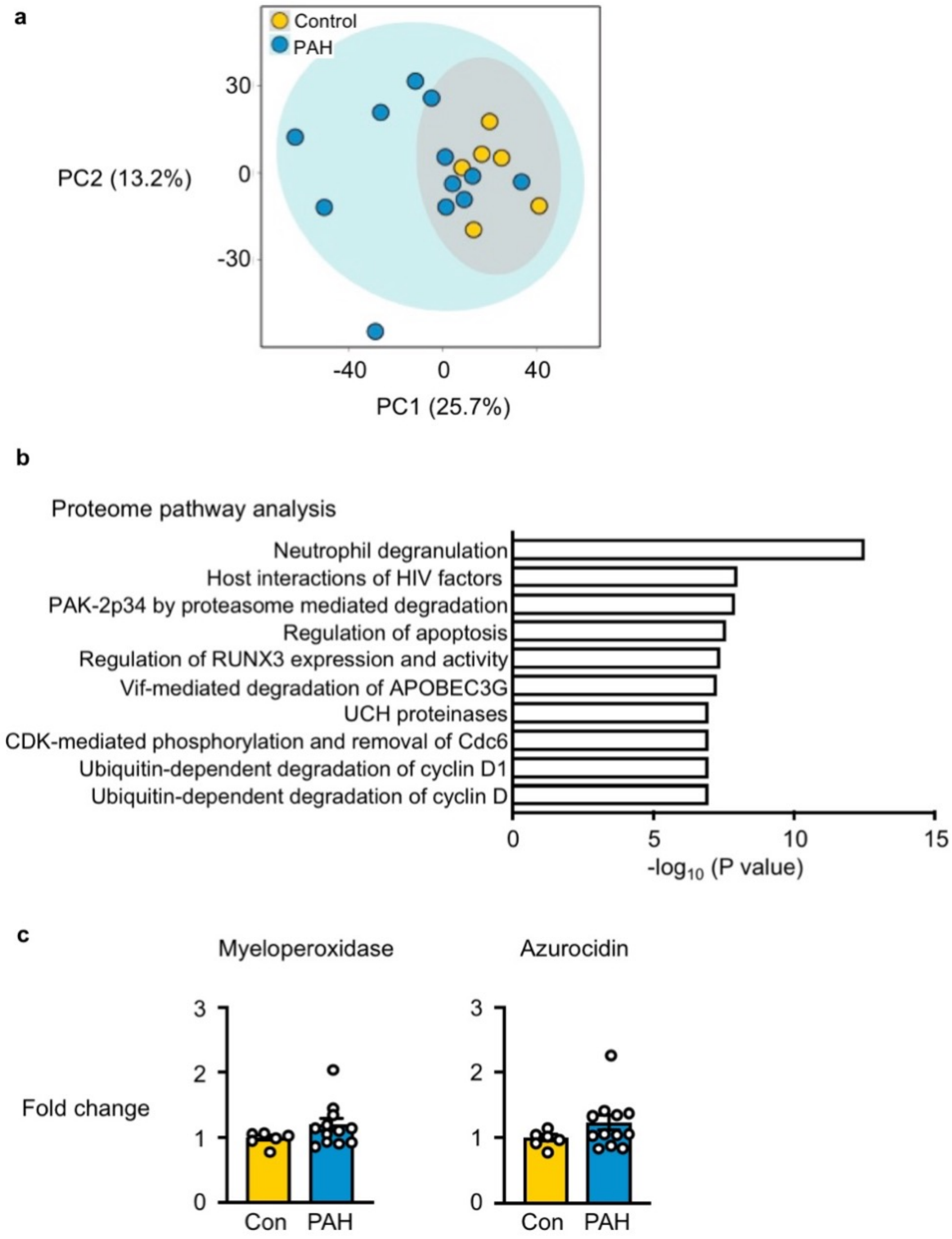


**Supplementary Figure 3. Identification of protein expression and pathways**

**(a)** Principal component analysis (PCA) plot of proteomics data that characterize the trends exhibited by the expression profiles of isolated PAH vs. Con neutrophils. Each dot represents a Con or PAH patient. **(b)** The top 10 pathways identified by Integrated Molecular Pathway-Level Analysis (IMPALA) of differentially expressed proteins in PAH vs. Con neutrophils. **(c)** Proteomic analysis of the fold-change in myeloperoxidase and Azurocidin in PAH vs. Con neutrophils (n=6 Con and n=12 PAH).

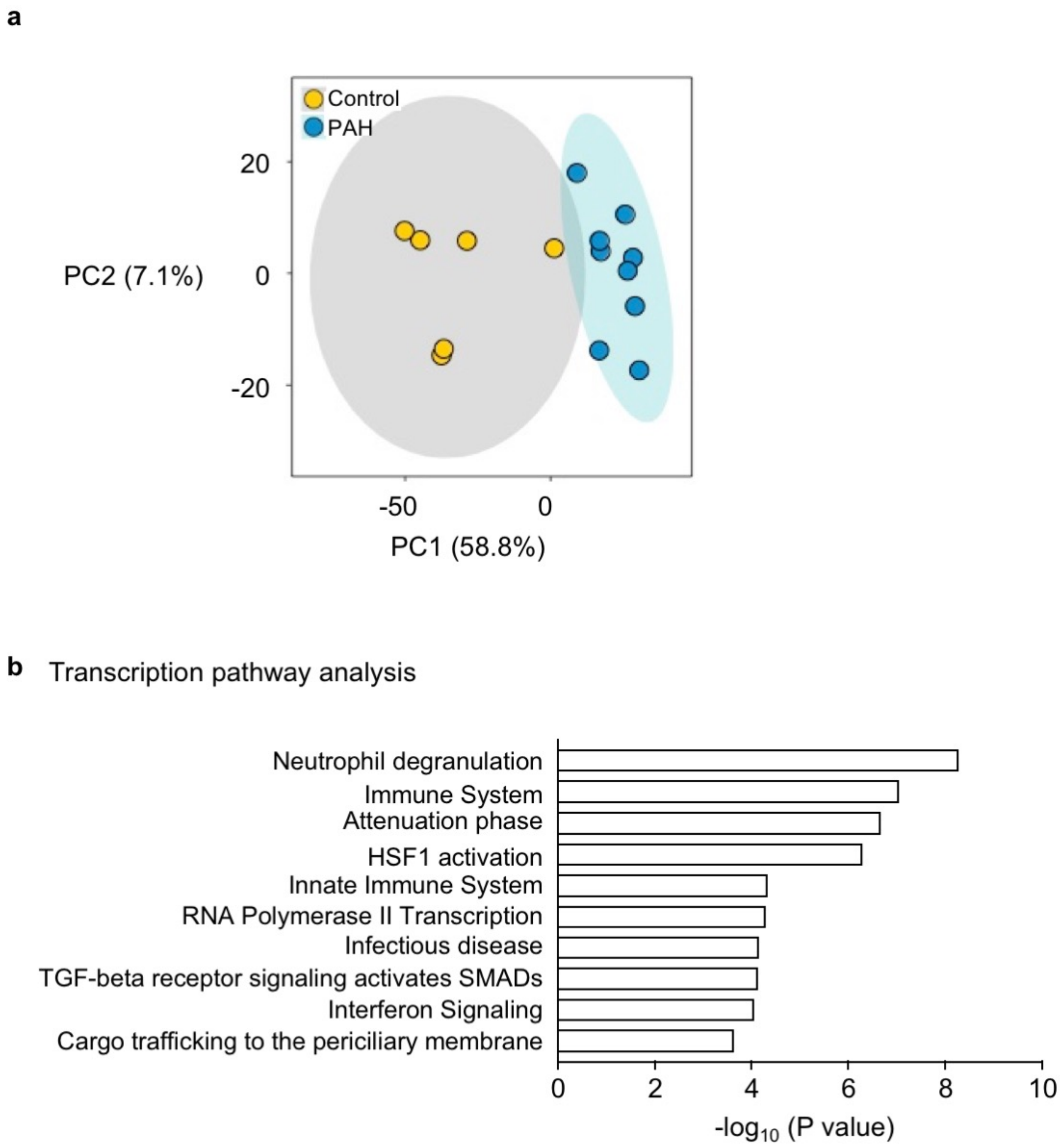


**Supplementary Figure 4. Identification of gene expression and pathways**

**(a)** Principal component analysis (PCA) plot of RNAseq data that characterizes the trends exhibited by the expression profiles of isolated PAH vs. Con neutrophils. Each dot represents a Con or PAH patient. **(b)** The top 10 pathways identified by Integrated Molecular Pathway-Level Analysis (IMPALA) of significant differentially expressed genes in PAH vs. Con neutrophils (n=6 Con and n=9 PAH patients).

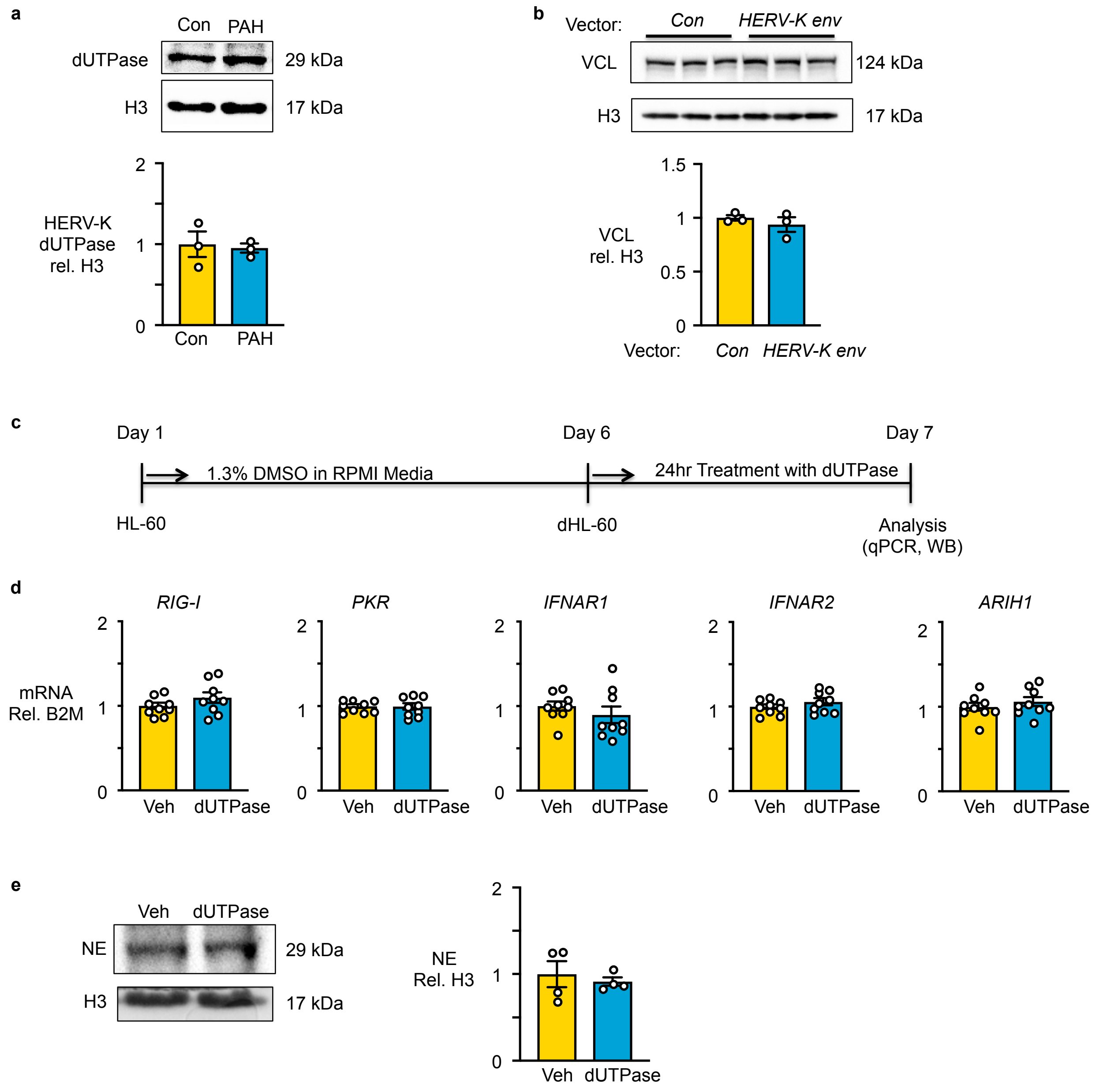


**Supplementary Figure 5. HERV dUTPase in neutrophils, and VCL and interferon following transfection of HL-60 cells with HERV-K envelope.**

**(a)** Representative western immunoblot and quantification of HERV-K dUTPase relative to histone H3 (H3) in PAH vs. Con neutrophils (n=3). **(b)** Representative western immunoblot and quantification of VCL relative to H3 in HL-60 cells overexpressing HERV-K envelope (n=3). **(c)** Illustration of the timeline of HL-60 cell differentiation and stimulation with recombinant HERV-K dUTPase**. (d)** PCR analysis of *RIG-I*, *PKR, IFNAR1, IFNAR2, and ARIH1* in dHL-60 cells treated with HERV-K dUTPase (dUTPase) or Vehicle (Veh; HERV-K dUTPase elution buffer) for 24 hr (n=9). **(e)** Representative western immunoblot and quantification of NE relative to H3 in dHL-60 cells treated with HERV-K dUTPase or Veh for 24 hr (n=4). Bars represent mean ± SEM.

### SUPPLEMENTARY TABLES

#### Supplementary Table 1. Demographics and assays used for PAH patients

| Sample | Gender | Age (Yr) | Race | Ethnicity | Diagnosis[1] | PAP[2] | PVR[3] | 6MWD[4] | NYHA[5] | Assays |
|---|---|---|---|---|---|---|---|---|---|---|
| PAH-1 | F | 30 | Other | Hispanic or Latino | IPAH | 43 | 4.8 | 525.79 | II | Transmigration Figure 2c (Sup 2a), TEM Figure 2d |
| PAH-2A | F | 37 | Other | Hispanic or Latino | IPAH | 20 | 2.7 | 502.93 | I | NE WB Figure 1a, NE Activity Extracellular Figure 1c, Transmigration Figure 2c (Sup 2a) |
| PAH-2B | F | 37 | Other | Hispanic or Latino | IPAH | 20 | 2.7 | 608.08 | I | NE activity Total lystate Figure 1b, Neutrophil/mL Figure 1d, NE activity/mL blood Figure 1e, NETosis Figure 1f, Proteomics Figure 3a-d, RNAseq Figure 4a-d |
| PAH-3 | F | 59 | White | Unknown | IPAH | 44 | 11.2 | 571.51 | I/II | Adhesion Figure 2a, TEM Figure 2d, dUTPase WB Figure 5e, Exosome Figure 6a-d, Animal Figure 7b-e |
| PAH-4A | F | 42 | Other | Hispanic or Latino | IPAH | 68 | 12.2 | 457.21 | II | dUTPase WB Figure 5e, Exosome Figure 6a-d, Animal Figure 7b-e |
| PAH-4B | F | 44 | Other | Hispanic or Latino | IPAH | 53 | 5.7 | 480.07 | II | HERV-K WB Figure 5a |
| PAH-5 | F | 31 | White | Unknown | IPAH | 48 | 11.4 | 557.79 | I | ITGB2 FACS Figure 3f, HERV-K WB Figure 5a, HERV-K Confocal Figure 5b, dUTPase WB Figure 5e, Exosome Figure 6a-d, Animal Figure 7b-e |
| PAH-6A | F | 47 | White | Not Hispanic or Latino | IPAH | 43 | 8.3 | 574.55 | I | PCR Figure 4e, dUTPase WB Figure 5e, Exosome Figure 6a-d, Animal Figure 7b-e |
| PAH-6B | F | 48 | White | Not Hispanic or Latino | IPAH | 43 | 8.3 | 594.37 | I | ITGB2 FACS Figure 3f, dUTPase WB Figure 5e, Exosome Figure 6b-e, Animal Figure 7b-e |
| PAH-7A | F | 65 | White | Not Hispanic or Latino | IPAH | 35 | 5.6 | 406.91 | II | NE WB Figure 1a, dUTPase WB Figure 5e, Exosome Figure 6a-d, Animal Figure 7b-e |
| PAH-7B | F | 66 | White | Not Hispanic or Latino | IPAH | 35 | 5.6 | 405.39 | II | VCL WB Figure 3e, ITGB2 FACS Figure 3f, HERV-K WB Figure 5a |
| PAH-8A | F | 38 | Asian | Not Hispanic or Latino | IPAH | 57 | 13.7 | 495.31 | II/III | Adhesion Figure 2a |
| PAH-8B | F | 48 | Asian | Not Hispanic or Latino | IPAH | 57 | 13.7 | 405.39 | III | NE WB Figure 1a, Chemokinesis Figure 2b, Proteomics Figure 3a-d, RNAseq Figure 4a-d |
| PAH-8C | F | 49 | Asian | Not Hispanic or Latino | IPAH | 57 | 13.7 | 585.22 | II | NE activity Total lystate Figure 1b, Neutrophil/mL Figure 1d, NE activity/mL blood Figure 1e, Transmigration Figure 2c (Sup 2a) |
| PAH-8D | F | 49 | Asian | Not Hispanic or Latino | IPAH | 58 | 12 | 548.65 | II/III | NETosis Figure 1f, HERV-K WB Figure 5a |
| PAH-8E | F | 50 | Asian | Not Hispanic or Latino | IPAH | 58 | 12 | 571.51 | II | VCL WB Figure 3e, dUTPase WB Figure 5e, Exosome Figure 6a-d, Animal Figure 7b-e |
| PAH-9A | F | 39 | Other | Hispanic or Latino | IPAH | 54 | 11.3 | 548.65 | II | NE WB Figure 1a, Adhesion Figure 2a, Transmigration Figure 2c (Sup 2a), Proteomics Figure 3a-d, RNAseq Figure 4a-d |
| PAH-9B | F | 40 | Other | Hispanic or Latino | IPAH | 66 | 13.8 | 548.65 | III | NE activity Total lystate Figure 1b, NE Activity Extracellular Figure 1c,Neutrophil/mL Figure 1d, NE activity/mL blood Figure 1e, Chemokinesis Figure 2b, dUTPase WB Figure 5e, Exosome Figure 6a-d, Animal Figure 7b-e |

| Sample | Gender | Age (Yr) | Race | Ethnicity | Diagnosis[1] | PAP[2] | PVR[3] | 6MWD[4] | NYHA[5] | Assays |
|---|---|---|---|---|---|---|---|---|---|---|
| PAH-10A | F | 35 | White | Not Hispanic or Latino | D&T associated PAH (anorexigen) Comorbidity: Mild Intermittent Asthma | 49 | 7.0 | 466.35 | II | NE WB Figure 1a |
| PAH-10B | F | 36 | White | Not Hispanic or Latino | D&T associated PAH (see PAH-10A) | 49 | 7.0 | 466.35 | III | NE WB Figure 1a, TEM Figure 2d |
| PAH-11 | F | 80 | White | Not Hispanic or Latino | IPAH | 52 | 17.7 | 292.61 | II | NE Activity Extracellular Figure 1c |
| PAH-12 | F | 73 | White | Not Hispanic or Latino | IPAH | 39 | 5.8 | 449.59 | II | NE activity Total lystate Figure 1b, Neutrophil/mL Figure 1d, NE activity/mL blood Figure 1e, TEM Figure 2d, Proteomics Figure 3a-d, RNAseq Figure 4a-d |
| PAH-13 | F | 52 | White | Hispanic or Latino | IPAH | 35 | 6.7 | 521.21 | II | NE WB Figure 1a, Proteomics Figure 3a-d, RNAseq Figure 4a-d |
| PAH-14A | F | 57 | White | Not Hispanic or Latino | IPAH | 62 | 16.5 | 384.05 | II/III | Adhesion Figure 2a |
| PAH-14B | F | 57 | White | Not Hispanic or Latino | IPAH | 62 | 16.5 | 411.49 | III | NE WB Figure 1a, Proteomics Figure 3a-d, RNAseq Figure 4a-d |
| PAH-14C | F | 58 | White | Not Hispanic or Latino | IPAH | 62 | 16.5 | 411.49 | II | NE activity Total lystate Figure 1b, Neutrophil/mL Figure 1d, NE activity/mL blood Figure 1e, NETosis Figure 1f |
| PAH-14D | F | 58 | White | Not Hispanic or Latino | IPAH | 62 | 16.5 | 429.77 | II | VCL WB Figure 3e, dUTPase WB Figure 5e, Exosome Figure 6a-d, Animal Figure 7b-e |
| PAH-14E | F | 59 | White | Not Hispanic or Latino | IPAH | 50 | 9.2 | 420.63 | II | ITGB2 FACS Figure 3f, HERV-K Confocal Figure 5b |
| PAH-14F | F | 60 | White | Not Hispanic or Latino | IPAH | 50 | 9.2 | 426.73 | II | HERV-K WB Figure 5a |
| PAH-15A | F | 50 | Black | Not Hispanic or Latino | IPAH | 52 | 7.6 | 361.19 | I/II | Adhesion Figure 2a, Chemokinesis Figure 2b |
| PAH-15B | F | 50 | Black | Not Hispanic or Latino | IPAH | 52 | 7.6 | 388.62 | I/II | NETosis Figure 1f |
| PAH-15C | F | 51 | Black | Not Hispanic or Latino | IPAH | 52 | 7.6 | 365.76 | II | Proteomics Figure 3a-d |
| PAH-16 | F | 53 | White | Not Hispanic or Latino | IPAH Comorbidity: COPD | 39 | 8 | 403.86 | II/III | NE WB Figure 1a, VCL WB Figure 3e, HERV-K WB Figure 5a, dUTPase WB Figure 5e, Exosome Figure 6a-d, Animal Figure 7b-e |
| PAH-17A | F | 61 | White | Not Hispanic or Latino | IPAH | 44 | 13.8 | 524.26 | III | NE activity Total lystate Figure 1b, Neutrophil/mL Figure 1d, NE activity/mL blood Figure 1e |
| PAH-17B | F | 62 | White | Not Hispanic or Latino | IPAH | 44 | 13.8 | 451.11 | III | TEM Figure 2d |
| PAH-17C | F | 63 | White | Not Hispanic or Latino | IPAH | 44 | 13.8 | 521.21 | III | PCR Figure 4e |

| Sample | Gender | Age (Yr) | Race | Ethnicity | Diagnosis[1] | PAP[2] | PVR[3] | 6MWD[4] | NYHA[5] | Assays |
|---|---|---|---|---|---|---|---|---|---|---|
| PAH-18 | F | 32 | White | Not Hispanic or Latino | IPAH | 45 | 10.6 | 734.58 | I | PCR Figure 4e, dUTPase WB Figure 5e, Exosome Figure 6a-d, Animal Figure 7b-e |
| PAH-19A | F | 65 | Other | Hispanic or Latino | IPAH | 60 | 11.1 | 365.76 | II | Adhesion Figure 2a |
| PAH-19B | F | 66 | Other | Hispanic or Latino | IPAH | 48 | 11 | 319.43 | III | Proteomics Figure 3a-d, RNAseq Figure 4a-d |
| PAH-19C | F | 66 | Other | Hispanic or Latino | IPAH | 48 | 11 | 374.91 | II | PCR Figure 4e |
| PAH-20 | F | 59 | Black | Not Hispanic or Latino | IPAH Comorbidity: Mild COPD | 61 | 17.1 | 315.47 | III | VCL WB Figure 3e, HERV-K WB Figure 5a |
| PAH-21 | F | 54 | Other | Not Hispanic or Latino | IPAH Comorbidity: End Stage Renal Disease | 41 | 6.1 | 274.32 | II/III | TEM Figure 2d, Proteomics Figure 3a-d, RNAseq Figure 4a-d |
| PAH-22A | F | 38 | Asian | Not Hispanic or Latino | IPAH | 33 | 3.8 | 624.85 | I | Proteomics Figure 3a-d |
| PAH-22B | F | 39 | Asian | Not Hispanic or Latino | IPAH | 33 | 3.8 | 617.23 | I | PCR Figure 4e |
| PAH-22C | F | 39 | Asian | Not Hispanic or Latino | IPAH | 33 | 3.8 | 594.37 | I | Adhesion Figure 2a |
| PAH-23A | F | 66 | Black | Not Hispanic or Latino | IPAH Comorbidity: Mild COPD, Sleep Disordered Breathing | 56 | 14.5 | 39.62 | III | Adhesion Figure 2a |
| PAH-23B | F | 68 | Black | Not Hispanic or Latino | IPAH Comorbidity: See PAH-23A | 69 | 15.6 | 73.15 | III | TEM Figure 2d, ITGB2 FACS Figure 3f |
| PAH-24 | F | 23 | White | Hispanic or Latino | IPAH | 33 | 6.8 | 551.09 | II | NE WB Figure 1a, NE activity Total lystate Figure 1b, Neutrophil/mL Figure 1d, NE activity/mL blood Figure 1e |
| PAH-25 | F | 29 | White | Not Hispanic or Latino | IPAH | 58 | 11.1 | 67.06 | IV | NE activity Total lystate Figure 1b, Neutrophil/mL Figure 1d, NE activity/mL blood Figure 1e |
| PAH-26 | F | 33 | Asian | Not Hispanic or Latino | IPAH | 51 | 8.1 | 525.79 | I/II | TEM Figure 2d |
| PAH-27A | F | 40 | White | Not Hispanic or Latino | IPAH | 25 | 5.4 | 585.22 | III | NE WB Figure 1a |
| PAH-27B | F | 40 | White | Not Hispanic or Latino | IPAH | 25 | 5.4 | 594.37 | II | NE activity Total lystate Figure 1b, Neutrophil/mL Figure 1d, NE activity/mL blood Figure 1e |
| PAH-27C | F | 40 | White | Not Hispanic or Latino | IPAH | 25 | 5.4 | 594.37 | II | ITGB2 FACS Figure 3f |
| PAH-27D | F | 42 | White | Not Hispanic or Latino | IPAH | 34 | 8.2 | 571.51 | II/III | dUTPase WB Figure 5e, Exosome Figure 6a-d, Animal Figure 7b-e |
| PAH-27E | F | 42 | White | Not Hispanic or Latino | IPAH | 34 | 8.2 | 594.37 | II/III | PCR Figure 4e, HERV-K WB Figure 5a, HERV-K Confocal Figure 5b |
| PAH-28 | F | 45 | White | Unknown | D&T assaciated PAH (Methamph etamine) | 52 | 20 | 658.38 | I/II | dUTPase WB Figure 5e, Exosome Figure 6a-d, Animal Figure 7b-e |

| Sample | Gender | Age (Yr) | Race | Ethnicity | Diagnosis[1] | PAP[2] | PVR[3] | 6MWD[4] | NYHA[5] | Assays |
|---|---|---|---|---|---|---|---|---|---|---|
| PAH-29 | F | 51 | Asian | Not Hispanic or Latino | IPAH | 35 | 7.8 | | II | VCL WB Figure 3e, HERV-K WB Figure 5a, dUTPase WB Figure 5e, Exosome Figure 6a-d, Animal Figure 7b-e |
| PAH-30A | F | 54 | Unkno wn | Unknown | IPAH | 48 | 12.2 | 556.27 | II | dUTPase WB Figure 5e, Exosome Figure 6a-d, Animal Figure 7b-e |
| PAH-30B | F | 54 | Unkno wn | Unknown | IPAH | 48 | 12.2 | 502.93 | II | PCR Figure 4e, HERV-K WB Figure 5a |
| PAH-31 | F | 49 | White | Not Hispanic or Latino | IPAH | 47 | 8 | 416.06 | III | PCR Figure 4e |
| PAH-32 | F | 51 | White | Not Hispanic or Latino | IPAH | 53 | 11.4 | 457.21 | II | ITGB2 FACS Figure 3f |
| PAH-33A | F | 33 | Other | Hispanic or Latino | IPAH | 58 | 14.7 | 441.97 | II | NE WB Figure 1a |
| PAH-33B | F | 33 | Other | Hispanic or Latino | IPAH | 58 | 14.7 | 310.90 | III | PCR Figure 4e |
| PAH-34A | F | 53 | White | Not Hispanic or Latino | IPAH | 33 | 5.8 | 566.93 | II | TEM Figure 2d |
| PAH-34B | F | 54 | White | Not Hispanic or Latino | IPAH | 33 | 5.8 | 603.51 | II | ITGB2 FACS Figure 3f, dUTPase WB Figure 5e, Exosome Figure 6a-d, Animal Figure 7b-e |
| PAH-35A | F | 45 | Black | Not Hispanic or Latino | IPAH | 38 | 6.1 | | II | NE activity Total lystate Figure 1b, Neutrophil/mL Figure 1d, NE activity/mL blood Figure 1, dUTPase WB Figure 5e Exosome Figure 6b-e, Animal Figure 7b-e |
| PAH-35B | F | 45 | Black | Not Hispanic or Latino | IPAH | 38 | 6.1 | | II | Proteomics Figure 3a-d, RNAseq Figure 4a-d |
| PAH-36 | F | 29 | White | Unknown | IPAH | 51 | 12.5 | 501.40 | I/II | Adhesion Figure 2a, Proteomics Figure 3a-d |

[1]Diagnosis: IPAH, Idiopathic PAH; D&T Drug and toxin associated PAH, COPD, Chronic obstructive pulmonary disease

[2]PAP: Mean Pulmonary Arterial Pressure, test closest to the time of blood draw

[3]PVR: Pulmonary vascular resistance in Woods Units, test closest to the time of blood draw

[4]6MWD: Distance (in meters) walked in six minutes, test closest to the time of blood draw

[5]NYHA: New York Heart Association functional classification

**Supplementary Table 2. Medication prescribed for PAH patients**

| Patient | PH medications |
|---|---|
| PAH-1 | sildenafil |
| PAH-2A | none |
| PAH-2B | none |
| PAH-3 | ambrisentan |
| PAH-4A | ambrisentan, sildenafil, treprostinil |
| PAH-4B | ambrisentan, sildenafil, treprostinil |
| PAH-5 | bosentan, sildenafil, treprostinil |
| PAH-6A | flolan |
| PAH-6B | epoprostenol |
| PAH-7A | macitentan, sildenafil |
| PAH-7B | macitentan, sildenafil |
| PAH-8A | tadalafil, flolan |
| PAH-8B | tadalafil, flolan |
| PAH-8C | tadalafil, flolan |
| PAH-8D | epoprostenol, tadalafil |
| PAH-8E | epoprostenol, tadalafil |
| PAH-9A | flolan, macitentan, sildenafil |
| PAH-9B | flolan, macitentan, sildenafil |
| PAH-10A | ambrisentan, tadalafil, treprostinil |
| PAH-10B | ambrisentan, tadalafil, remodulin |
| PAH-11 | macitentan |
| PAH-12 | macitentan, tadalafil |
| PAH-13 | ambrisentan, sildenafil, tyvaso |
| PAH-14A | ambrisentan, sildenafil, flolan |
| PAH-14B | ambrisentan, sildenafil, flolan |
| PAH-14C | ambrisentan, epoprostenol, sildenafil |
| PAH-14D | ambrisentan, epoprostenol, sildenafil |
| PAH-14E | ambrisentan, epoprostenol, sildenafil |
| PAH-14F | ambrisentan, epoprostenol, sildenafil |
| PAH-15A | tadalafil, treprostinil |
| PAH-15B | tadalafil, treprostinil |
| PAH-15C | tadalafil, treprostinil |
| PAH-16 | tadalafil |
| PAH-17A | bosentan, sildenafil |

| Patient | PH medications |
|---|---|
| PAH-17B | bosentan, selexipag, sildenafil |
| PAH-17C | bosentan, sildenafil |
| PAH-18 | macitentan, tadalafil |
| PAH-19A | ambrisentan, flolan |
| PAH-19B | ambrisentan, flolan |
| PAH-19C | ambrisentan, flolan |
| PAH-20 | sildenafil, treprostinil |
| PAH-21 | remodulin |
| PAH-22A | tadalafil, treprostinil |
| PAH-22B | tadalafil, treprostinil |
| PAH-22C | tadalafil, treprostinil |
| PAH-23A | sildenafil, treprostinil |
| PAH-23B | macitentan, sildenafil, remodulin |
| PAH-24 | ambrisentan, tadalafil, treprostinil |
| PAH-25 | ambrisentan, sildenafil, treprostinil |
| PAH-26 | remodulin, sildenafil |
| PAH-27A | sildenafil |
| PAH-27B | macitentan, sildenafil |
| PAH-27C | macitentan, sildenafil |
| PAH-27D | macitentan, sildenafil |
| PAH-27E | macitentan, sildenafil |
| PAH-28 | ambrisentan, tadalafil, treprostinil |
| PAH-29 | ambrisentan, sildenafil |
| PAH-30A | ambrisentan |
| PAH-30B | ambrisentan, tadalafil, treprostinil |
| PAH-31 | ambrisentan, selexipag, sildenafil |
| PAH-32 | macitentan, sildenafil |
| PAH-33A | ambrisentan, sildenafil, remodulin |
| PAH-33B | ambrisentan, sildenafil, remodulin |
| PAH-34A | bosentan, epoprostenol |
| PAH-34B | bosentan, epoprostenol |
| PAH-35A | iloprost, sildenafil |
| PAH-35B | iloprost, sildenafil |
| PAH-36 | tadalafil, flolan |

**Supplementary Table 3. Demographics of Donor Controls, and Assays**

| Sample | Age (Yr) | Gender | Race | Ethnicity | Echo done (Yes/No) | IVS | PW | LVID | LVEF | E/e’ | Estimated RAP | Assays |
|---|---|---|---|---|---|---|---|---|---|---|---|---|
| CON-1 | 30 | M | White | Hispanic | No | | | | | | | NE activity Total lystate Figure 1b, NETosis Figure 1d, RNAseq Figure 4a-d |
| CON-2 | 29 | F | White | Non-Hispanic | Yes | 0.78 | 0.76 | 4.3 | 64.5 | 5.3 | 5 | Adhesion Figure 2a |
| CON-3 | 41 | F | Asian | Non-Hispanic | Yes | 0.7 | 0.71 | 4.4 | 61 | 6.9 | 5 | NE WB Figure 1a, NETosis Figure 1d, Adhesion Figure 2a, Proteomics Figure 3a-d, VCL WB Figure 3e, RNAseq Figure 4a-d, HERV-K WB Figure 5a, HERV-K Confocal Figure 5b, dUTPase WB Figure 5e, Exosome Figure 6a-d, Animal Figure 7b-e |
| CON-4 | 29 | F | Caucasian | Non-Hispanic | Yes | | | | | | | NE WB Figure 1a, NETosis Figure 1d, Proteomics Figure 3a-d, RNAseq Figure 4a-d |
| CON-5 | 34 | F | Asian | Non-Hispanic | No | | | | | | | NETosis Figure 1d, VCL WB Figure 3e, ITGB2 FACS Figure 3f, PCR Figure 4e, HERV-K WB Figure 5a, HERV-K Confocal Figure 5b, dUTPase WB Figure 5e, Exosome Figure 6a-d, Animal Figure 7b-e |
| CON-6 | 49 | F | White | Hispanic | Yes | 0.86 | 0.86 | 4.9 | 63.3 | 10.1 | 5 | NE activity Total lystate Figure 1b, NETosis Figure 1d, Proteomics Figure 3a-d |
| CON-7 | 32 | F | American Indian/Alaska Native | Hispanic | Yes | 0.94 | 0.87 | 4.4 | 65.6 | 8.2 | 5 | Adhesion Figure 2a, Proteomics Figure 3a-d, RNAseq Figure 4a-d |
| CON-8 | 41 | F | Black | Non-Hispanic | Yes | 0.75 | 0.69 | 4.1 | 69.9 | 8.6 | 5 | Proteomics Figure 3a-d, RNAseq Figure 4a-d |
| CON-9 | 50 | F | Black | Non-Hispanic | Yes | 1.3 | 1.3 | 5.3 | 64.8 | 11.2 | 5 | NE activity Total lystate Figure 1b, Proteomics Figure 3a-d, RNAseq Figure 4a-d |
| CON-10 | 33 | F | Black | Non-Hispanic | Yes | 0.69 | 0.77 | 4.6 | 68.4 | 8.3 | 5 | VCL WB Figure 3e, ITGB2 FACS Figure 3f, dUTPase WB Figure5 e, Exosome Figure 6a-d, Animal Figure 7b-e |
| CON-11 | 40 | F | White | Non-Hispanic | No | | | | | | | VCL WB Figure 3e, PCR Figure 4e, HERV-K WB Figure 5a, HERV-K Confocal Figure 5b, dUTPase WB Figure 5e, Exosome Figure 6a-d, Animal Figure 7b-e |
| CON-12 | 40 | M | White | Non-Hispanic | No | | | | | | | NE WB Figure 1a, VCL WB Figure 3e, ITGB2 FACS Figure 3f, HERV-K WB Figure 5a, dUTPase WB Figure 5e, Exosome Figure 6a-d, Animal Figure 7b-e |
| CON-13 | 35 | F | White | Non-Hispanic | No | | | | | | | NETosis Figure 1d, dUTPase WB Figure 5e, Exosome Figure 6a-d, Animal Figure 7b-e |
| CON-14 | 24 | F | White | Non-Hispanic | No | | | | | | | ITGB2 FACS Figure 3f, dUTPase WB Figure 5e, Exosome Figure 6a-d, Animal Figure 7b-e |
| CON-15 | 36 | F | Asian | Non-Hispanic | No | | | | | | | dUTPase WB Figure5e, Exosome Figure 6a-d, Animal Figure 7b-e |
| CON-16 | 31 | F | Not reported | Not reported | No | | | | | | | dUTPase WB Figure5e, Exosome Figure 6a-d, Animal Figure 7b-e |
| CON-17 | 31 | F | White | Non-Hispanic | No | | | | | | | PCR Figure 4e, dUTPase WB Figure5e, Exosome Figure 6a-d, Animal Figure 7b-e |
| CON-18 | 60 | F | Black | Non-Hispanic | No | | | | | | | PCR Figure 4e, dUTPase WB Figure5e, Exosome Figure 6a-d, Animal Figure 7b-e |

Age (years); IVS: interventricular septum (cm); LVEF: left ventricular ejection fraction (%); LVID: left ventricular internal dimension (cm); PW: posterior wall (cm); RAP: right atrial pressure (mmHg); E/e’: ratio between early mitral inflow velocity and mitral annular early diastolic velocity.

**Supplementary Table 4. PAEC Demographics**

**a. Donor controls**

| Patient | Age (yr) | Gender | Race | Ethnicity | Cause of Death |
|---|---|---|---|---|---|
| EC CON1 | 33 | F | White | Non-Hispanic | Head trauma. Blunt injury. |
| EC CON2 | 57 | F | White | Non-Hispanic | Acute Myocardial Infarction |
| EC CON3 | 1 | M | White | Non-Hispanic | Anoxia/Drowning |

**b. PAH Patients**

| Patient | Age (yr) | Gender | Race | Ethnicity | Diagnosis[1] | PAP[2] | PVR[3] | 6MWD[4] | PAH Meds |
|---|---|---|---|---|---|---|---|---|---|
| EC PAH1 | 22 | F | White | Non-Hispanic | FPAH | 82 | 16.04 | 375.82 | sildenafil, ambrisentan, tadalafil, inhaled treprostinil, treprostinil (orenitram) |
| EC PAH2 | 27 | F | White | Non-Hispanic | IPAH | 69 | 12.11 | 359.66 | sildenafil, SC treprostinil, IV treprostinil, bosentan, inhaled iloprost |
| EC PAH3 | 53 | M | White | Non-Hispanic | IPAH | 33 | 3.86 | 77.72 | sildenafil, tadalafil, macitentan, IV epoprostenol |

[1]Diagnosis: FPAH, Familial PAH; IPAH, Idiopathic PAH

[2]PAP: Mean Pulmonary Arterial Pressure, test closest to the time of blood draw

[3]PVR: Pulmonary vascular resistance in Woods Units, test closest to the time of blood draw

[4]6MWD: Distance (in meters) walked in six minutes, test closest to the time of blood draw

**Supplementary Table 5. Correlative Genes and Proteins from the Omics Analysis**

| Gene/Protein | Correlation | Protein p-value | mRNA p-value |
|---|---|---|---|
| EIF3J | Decreased | 0.003299398 | 0.000164377 |
| TUBB4B | Decreased | 0.019693895 | 0.0000112 |
| TCP1 | Decreased | 0.001135572 | 0.0000459 |
| SUN2 | Decreased | 0.005046265 | 0.0000971 |
| TPD52L2 | Decreased | 0.000716389 | 0.00000437 |
| PCYT2 | Decreased | 0.008718198 | 0.000219796 |
| STOML2 | Decreased | 0.016078615 | 0.000000427 |
| PSMC4 | Decreased | 0.013149119 | 0.00000294 |
| EEF1D | Decreased | 0.006689322 | 0.000320571 |
| MECP2 | Decreased | 0.009342118 | 0.0000423 |
| POLR2A | Decreased | 0.018024105 | 0.002416794 |
| RALY | Decreased | 0.009951085 | 0.003563528 |
| SF1 | Decreased | 0.014691391 | 0.000407631 |
| AMPD2 | Decreased | 0.000421312 | 0.000422481 |
| CHD4 | Decreased | 0.023893101 | 4.52E-09 |
| SRRT | Decreased | 0.01383299 | 0.0000182 |
| ARIH1 | Decreased | 0.032208307 | 0.00001 |
| COPS5 | Increased | 0.013280254 | 0.004345779 |
| VBP1 | Increased | 0.032490686 | 0.00188536 |
| VPS36 | Increased | 0.021535145 | 0.002750826 |
| PFN1 | Increased | 0.014676215 | 0.002221079 |
| CSTA | Increased | 0.003669109 | 0.000904283 |
| TM9SF2 | Increased | 0.005699783 | 0.0000727 |
| PGGT1B | Increased | 0.027796704 | 0.000186515 |
| PGAM1 | Increased | 0.030360717 | 0.001498589 |
| BROX | Increased | 0.010059018 | 0.001607269 |
| ARCN1 | Increased | 0.000267714 | 0.001629735 |
| MYCBP | Increased | 0.029877858 | 0.001891447 |
| SH3BGRL | Increased | 0.001591546 | 0.001628348 |
| UQCR10 | Increased | 0.006935534 | 0.000919845 |
| CTSS | Increased | 0.006901583 | 0.002102924 |
| MTARC1 | Increased | 0.001422351 | 0.000791293 |
| SELL | Increased | 0.012844352 | 0.00010617 |

**Supplementary Table 6. Reagents and Resources**

| REAGENT or RESOURCE | SOURCE | IDENTIFIER/SEQUENCE |
|---|---|---|
| **Antibodies** | | |
| Anti-human CD66b/CEACAM 8 | Sigma | Cat#: SAB4301144-100 |
| Anti-Neutrophil Elastase | Abcam | Cat#: ab68672 |
| Anti-Histone H3 | Abcam | Cat#: ab24834 |
| Anti-Vinculin | Abcam | Cat#: ab18058 |
| Anti-HERV-K Envelope | Ango | Cat#: Herm 1811-5 |
| Anti-dUTPase | Dr. Ariza | Cat#: ab265 |
| CD9 Monoclonal Antibody (Ts9) | ThermoFisher | Cat #: 10626D |
| Anti-ITGB2 Flor antibody | Novus | Cat#: NB500-480AF700 |
| GAPDH Antibody (G-9) | Santa Cruz | Cat#: sc-365062 |
| Alexa Fluor488 donkey anti mouse IgG (H +L) | ThermoFisher | Cat#: A21202 |
| α-smooth muscle actin antibody | Sigma-Aldrich | Cat#: A2547 |
| CD9 Antibody | ThermoFisher | Cat#: 10626D |
| **Vectors** | | |
| EGFP control vector | VectorBuilder | ID: VB180814-1915cef |
| HERV-K Env vector | VectorBuilder | ID: VB180814-1914ntz |
| **Reagents, Cells, Chemicals, Peptides, and Recombinant Proteins** | | |
| HL-60 Cell Avalanche™ Transfection Reagent | EZ biosystems | Cat#: EZT-HL60-1 |
| RPMI 1640 | Life Technologies | Cat#: 11875119 |
| GIBCO | Life Technologies | Cat#: 10438026 |
| P/S | Life Technologies | Cat#: 15140122 |
| DMSO | Sigma Aldrich | Cat#: D2650-100ML |
| Sytox green nucleic acid | ThermoFisher | Cat# S7020 |
| DAPI Fluoromount-G | SouthernBiotech | Cat#: 0100-20 |
| Fibronectin from human plasma | Sigma Aldrich | Cat#: F0895-2MG |

| REAGENT or RESOURCE | SOURCE | IDENTIFIER/SEQUENCE |
|---|---|---|
| Recombinant Human CXCL8/IL-8 | R&D Systems | Cat#: 208-IL-050 |
| Elafin | Proteo-Biotech-AG | Gift |
| Calcein AM, Cell Permeant Dye | ThermoScientific | Cat#: C3100MP |
| DPBS | ThermoFisher | Cat#: 10010023 |
| Poly D lysine Solution | EMD Millipore | Cat#: A-003-E |
| N-Formyl-Met-Leu-Phe | Sigma Aldrich | Cat#: F3506-10MG |
| 1M Tris-HCL | Life Technologies | Cat#: 15567-027 |
| UltraPure 10%SDS | Life Technologies | Cat#: 24730-020 |
| HL-60 | Sigma Aldrich | Cat#: 98070106-1VL |
| Bovine Serum Albumin | Sigma Aldrich | Cat#: A3059-500G |
| Glycine 1 M solution | Millipore | Cat#: 67419-1ML-F |
| Formvar/Carbon Film | Electron Microscopy Science | Cat#: FCF300-CU |
| **Commercial Assays and Kits** | | |
| Dynabeads Antibody Coupling Kit 14311D | ThermoFisher (Invitrogen) | Cat#: 14311D |
| EnzChek™ Elastase Assay Kit | Thermofisher Scientific | Cat#: E12056 |
| MACSxpress Neutrophil Isolation Kit | MACS Miltenyi Biotec | Cat#: 130-104-434 |
| SMARTer® Stranded Total RNA-Seq Kit - Pico Input Mammalian - 12 Rxns | TAKARA | Cat#: 635005 |
| Albumin depletion kit | Thermofisher scientific | Cat#: 85160 |
| Corning® FluoroBlok™ Inserts, 24 well | Corning Incorporated Life Sciences | Cat#: 351151 |
| Direct-zol RNA Kits | Zymoresearch | Cat#: R2062 |
| DakoArk | Dako | Cat#: K3954 |
| Dako Liquid DAB + Substrate Chromogen System | Dako | Cat#: K3468 |

**Supplementary Table 7. Primers used for qPCR**

| GENE | SOURCE | FORWARD | REVERSE |
|---|---|---|---|
| PKR | Stanford PAN facility | tggaaagcgaacaaggagtaag | ccaaagcgtagaggtccactt |
| RIG-I | Stanford PAN facility | tgtgggcaatgtcatcaaaa | gaagcacttgctacctcttgc |
| IFNAR2 | Stanford PAN facility | ccttaaaatgcaccctccttc | ttcctcctattttggcagattc |
| IFNAR1 | Stanford PAN facility | aagggcgaggacgaagag | tcgacctctactttttgaggaga |
| IFNGR2 | Stanford PAN facility | gctgctgctcggagtctt | ggtctccaaagttagctgtgct |
| ARIH1 | Stanford PAN facility | tgggataaagagaagctaatgga | attaattacatgacactcagcaaagag |
| HERV-K env | Applied Biosystems | gctgccctgccaaacctgag | cctgagtgacatcccgcttacc |
| B2M | Stanford PAN facility | ttctggcctggaggctatc | tcaggaaatttgactttccattc |

**SUPPLEMENTARY VIDEO**

**Supplementary Video 1. Time Lapse imaging demonstration reduced migration (chemokinesis) in PAH vs. Control neutrophils in response to IL-8 (100 ng/mL).** Migration was observed using a confocal laser-scanning microscope (FV1000, Olympus, Center Valley, PA) with a 40X objective. Frames were taken every 10 sec for 30 min and analyzed using ImageJ software. Graphic analysis of several experiments is presented in Figure 2b.
